# In depth exploration of the alternative proteome of *Drosophila melanogaster*

**DOI:** 10.1101/2022.03.16.484563

**Authors:** Bertrand Fabre, Sebastien A Choteau, Carine Duboé, Carole Pichereaux, Audrey Montigny, Dagmara Korona, Michael J. Deery, Mylène Camus, Christine Brun, Odile Burlet-Schiltz, Steven Russell, Jean-Philippe Combier, Kathryn S Lilley, Serge Plaza

**Author notes:** Corresponding author: Fabre, Bertrand and Plaza, Serge.

## Abstract

Recent studies have shown that hundreds of small proteins were occulted when protein coding genes were annotated. These proteins, called alternative proteins, have failed to be annotated notably due to the short length of their open reading frame (less than 100 codons) or the enforced rule establishing that messenger RNAs (mRNAs) are monocystronic. Several alternative proteins were shown to be biologically active molecules and seem to be involved in a wide range of biological functions. However, genome wide exploration of the alternative proteome is still limited to a few species. In the present article we describe a deep peptidomics workflow which enabled the identification of 401 alternative proteins in *Drosophila melanogaster*. Subcellular localization, protein domains and short linear motifs were predicted for 235 of the alternative proteins identified and point towards specific functions of these small proteins. Several alternative proteins had approximated abundances higher than their canonical counterparts, suggesting that these alternative proteins are actually the main products of their corresponding genes. Finally, we observed fourteen alternative proteins with developmentally regulated expression patterns and ten induced upon heat shock treatment of embryos, demonstrating stage or stress specific production of alternative proteins.

## Introduction

Almost twenty years after the completion of the sequencing of the genomes of *Saccharomyces cerevisiae*, *Drosophila melanogaster* and *Homo sapiens*, precise gene annotation still remains challenging. Initiatives such as the Human Proteome Project (HPP) (Omenn et al., 2021) or ProteomicsDB (Lautenbacher et al., 2021) aim at defining the ensemble of proteins actually expressed in human or other organisms using mass spectrometry (MS) based approaches. These projects have reached impressive milestones but they are centered around the protein database that are used to mine the experimental MS data in order to identify expressed proteins (Brunet et al., 2020). So far, these databases mainly comprise genes annotated in UniProt (Bateman et al., 2021). However, recent studies have suggested that hundreds of small yet to be annotated proteins, might be expressed across the kingdom of life (Fabre et al., 2021). These proteins, called alternative proteins (AltProts, or short open reading frame (ORF) encoding polypeptides (SEPs) or microproteins), have failed to be annotated notably due to the short length of their open reading frame (less than 100 codons), alternative start codon (other than AUG) or the enforced rule establishing that messenger RNAs (mRNAs) are monocystronic (Brunet et al., 2020). Almost two decades of pioneering work have highlighted that AltProts can be produced from ORFs on long non-coding RNA (lncRNA-ORF) or the different regions of mRNAs, within the 5’ or 3’ untranslated regions or alternative frames in canonical coding sequences (called uORFs, dORFs and intORFs, respectively) (Plaza et al., 2017a). Database such as OpenProt (Brunet et al., 2019), https://sORFs.org (Olexiouk et al., 2018), SmProt (Li et al., 2021), ARA-PEPs (Hazarika et al., 2017), PsORF (Chen et al., 2020b) or MetamORF (Choteau et al., 2021) constitute repositories predicting the existence of potentially thousands of AltProts based mainly on ribosome footprints determined via ribosome profiling experiments. However, in most cases, we still lack unambiguous empirical evidence for the existence of most of these predicted short proteins. Although ribosome profiling approaches clearly established the binding of ribosome to alternative ORFs, it is in fact difficult to deduce the productive translation of the ORFs, resulting in the expression of stable proteins (Patraquim et al., 2020). Mass spectrometry is generally the method of choice for large scale identification of proteins and peptides (Cassidy et al., 2021). MS data demonstrating the genome wide expression of AltProts is still limited to few species (Fabre et al., 2021). The roles of only few alternative proteins, less than 50 across all species, have been characterized to date (Plaza et al., 2017b; Wright et al., 2021). The alternative proteins, which function has been determined, seem to be involved in a wide range of key biological processes (Plaza et al., 2017b; Wright et al., 2021). Due to their large spectrum of functions, alternative proteins represent an attractive new repertoire of molecules for drug development and agricultural applications.

In an effort to assess the pervasive production of alternative proteins in the model organism *Drosophila melanogaster*, we describe here the development of a deep peptidomics workflow combining different protein extraction methods, small protein enrichment steps, state of the art mass spectrometry and optimized bioinformatics analysis using the well curated OpenProt database. We were able to identify 401 yet unannotated alternative proteins, substantially increasing (twice) the repertoire of alternative proteins in *Drosophila melanogaster*. The majority of these proteins are produced from alternative reading frames in the canonical coding sequences (CDS), highlighting the fact that the proteome is more complex than previously anticipated. AltProts produced from different type of RNA (lncRNA or mRNA) or different regions of mRNA (5’ or 3’ untranslated regions or alternative frames within canonical CDS) have different amino acid compositions, isoelectric points, predicted protein domains or disordered regions. Surprisingly, AltProts are predicted to be localized mainly in cell nucleus, mitochondria or secreted. We identified several AltProts for which the approximated abundances were higher than their canonical counterparts, suggesting that these AltProts are actually the main products of their corresponding genes. Finally, we observed fourteen AltProts with developmentally regulated expression patterns and ten induced upon heat shock treatment of embryos, demonstrating stage or stress specific production of alternative proteins.

## Material and Methods

### Drosophila collection and S2 cell culture

*D. melanogaster* adult flies and embryos were maintained and collected as previously described (Fabre et al., 2019). S2 cells were cultured as described in (Montigny et al., 2021).

### Protein extraction and alternative protein enrichment

Several approaches were used to extract and enrich alternative proteins:

1. Embryo (100 μl equivalent of embryo per replicate), adult flies (10 adult flies per replicate) or S2 cell pellets (5 × 10^8^ cells per replicate) were resuspended in an SDS buffer (Tris 50 mM pH 7.5, 5% SDS), then immediately sonicated and boiled for 10 min at 95°C. A detergent compatible protein assay (Bio-Rad) was used to measure protein concentration. Loading buffer (Tris 40 mM pH 7.5, 2% SDS, 10% glycerol, 25 mM DTT final concentration) was added to 100 μg of protein per condition and samples were boiled for 5 min at 95°C. Proteins were alkylated using chloroacetamide at a final concentration of 60 mM for 30 min at room temperature in the dark. Samples were loaded on an SDS-PAGE gel (acrylamide concentration of 4% for the stacking gel and 12% for the resolving gel). After protein migration, staining with InstantBlue™ (Merck) was performed and bands were excised between 15 kDa and the dye front (3 bands for S2 cells and 2 bands for embryos and adult flies). Proteins were then digested over night at 37°C with trypsin (or glu-C or chymotrypsin in the case of S2 cells) using in gel digestion as previously described (Fabre et al., 2016b). Resulting peptides were injected on a Thermofisher Q Exactive plus (S2 cells samples only) or a Thermofisher Fusion (embryo and adult flies samples only). Three biological replicates were performed for each condition.
2. Embryo (200 μl equivalent of embryo per replicate) and adult flies (50 adult flies per replicate) were lysed and proteins were reduced and alkylated as described in the approach 1 and 1 mg of protein were digested using in gel digestion (trypsin for adult flies, or trypsin, Glu-C or chymotrypsin for embryos). The resulting peptides were then separated by high pH reverse phase fractionation as described in (Fabre et al., 2017). Each fraction was analyzed either on a Sciex TripleTOF 6600 (both embryo and adult flies samples), a Thermofisher Q Exactive (embryo samples only) or a Thermofisher Fusion Lumos (embryo samples only). Three biological replicates were performed for each condition.
3. Embryos (100 μl equivalent of embryo per replicate) were incubated at 37 °C to induce heat-shock or maintained at 25 °C as described previously (Fabre et al., 2016c). Proteins were extracted, reduced and alkylated as described in protocol 1 and 100 μg were loaded on an SDS-PAGE gel (acrylamide concentration of 4% for the stacking gel and 12% for the resolving gel). After a short migration each gel lane was cut in three bands and in gel digestion was performed with trypsin as previously described (Fabre et al., 2016b). Resulting peptides were injected on a Thermofisher Q Exactive. Three biological replicates were performed for each condition.
4. Embryos (100 μl equivalent of embryo per replicate) staged every 4.5 hours as previously described (Fabre et al., 2016a) were lysed in a buffer containing 20 mM HEPES pH 8, 150 mM KCl and 10 mM MgCl_2_ and proteins were first digested with proteinase K and boiled for 10 min after addition of Guanidine hydrochloride (GnHCl) at a 6 M final concentration. The proteins were then reduced with 25 mM dithiothreitol (DTT), alkylated with chloroacetamide at a final concentration of 60 mM and digested with trypsin, glu-C or chymotrypsin over night at 37°C. Peptides were desalted on a C18 SepPak column (Waters), dried down using a speed-vac, labelled with Tandem Mass Tag (TMT) 10-plex (Thermo Scientific) according to the manufacturer’s instructions, pooled and fractionated using the High pH Reversed-Phase Peptide Fractionation Kit (Pierce). Each fraction was analyzed on a Thermofisher Fusion Lumos. Three biological replicates were performed for each condition.
5. 5 × 10^8^ S2 cells were boiled at 95°C for 20 min in water and sonicated. Then acetic acid and acetonitrile were added to the sample both at a final concentration of 20% and 5% respectively. Samples were centrifuged at 20,000 g for 20 min at 4°C and the pellet was discarded. Supernatant were dried using a speed-vac and proteins were resuspended in 6 M GnHCl, 50 mM ammonium bicarbonate. A BCA assay (Pierce) was used to measure protein concentration. Proteins were reduced in 5 mM TCEP (tris 2-carboxyethylphosphine hydrochloride) for 1 h at 37 °C and alkylated in 10 mM chloroacetamide for 30 min at RT in the dark. The samples were diluted with 50 mM ammonium bicarbonate to a final concentration of GnHCl of 0.5 M. Proteins were digested with trypsin (at a 1:50 trypsin to protein ratio) and resulting peptides were desalted on a C18 Hypersep column (Thermo Scientific) and dried down using a speed-vac. Samples were injected on a Thermofisher Fusion. Two biological replicates were performed.
6. 5 × 10^8^ S2 cells were boiled at 95°C for 20 min in GnHCl lysis buffer (6 M Guanidine Hydrochloryde, Tris 50 mM pH 7.5 and 100 mM NaCl) and sonicated. Samples were centrifuged at 20,000 g for 20 min at RT and the pellet was discarded. Trifluoroacetic acid (TFA) was added to the supernatant to a final concentration of 0.4% before loading the sample on a C8 column (Pierce) preconditioned with acetonitrile (ACN) and equilibrated with 0.1% TFA. Column were washed twice with 0.1% TFA and proteins were eluted with 75% ACN, 0.1% TFA. Samples were dried down using a speed-vac and resuspended in 6 M GnHCl in 50 mM ammonium bicarbonate. A BCA assay (Pierce) was used to measure protein concentration. Proteins were reduced in 5 mM TCEP (tris 2-carboxyethylphosphine hydrochloride) for 1 h at 37 °C and alkylated in 10 mM chloroacetamide for 30 min at RT in the dark. The samples were diluted with 50 mM ammonium bicarbonate to a final concentration of GnHCl of 0.5 M. Proteins were digested with trypsin (at a 1:50 trypsin to protein ratio) and resulting peptides were desalted on a C18 Hypersep column (Thermo Scientific) and dried down using a speed-vac. Samples were injected on a Thermofisher Fusion. Three biological replicates were performed.
7. 5 × 10^8^ S2 cells were boiled at 95°C for 20 min in GnHCl lysis buffer (6 M Guanidine Hydrochloryde, Tris 50 mM pH 7.5 and 100 mM NaCl) and sonicated. The sample was centrifuged at 20,000 g for 20 min at RT and the pellet was discarded. The supernatant was loaded on an ultrafiltration device with a molecular weight cut-off of 30 kDa (Millipore) and the fraction retained (above 30 kDa) was discarded. A BCA assay (Pierce) was used to measure protein concentration. Proteins were reduced in 5 mM TCEP (tris 2-carboxyethylphosphine hydrochloride) for 1 h at 37 °C and alkylated in 10 mM chloroacetamide for 30 min at RT in the dark. The samples were diluted with 50 mM ammonium bicarbonate to a final concentration of GnHCl of 0.5 M. Proteins were digested with trypsin (at a 1:50 trypsin to protein ratio) and resulting peptides were desalted on a C18 Hypersep column (Thermo Scientific) and dried down using a speed-vac. Samples were injected on a Thermofisher Orbitrap Velos. One biological replicate was performed.

### Mass spectrometry analysis

Sciex TripleTOF 6600 and Thermofisher Q Exactive were operated as described in (Mata et al., 2017). The Thermofisher OrbiTrap Fusion Lumos was used as in (Geladaki et al., 2019). The Thermofisher OrbiTrap Velos and Q Exactive plus were operated as described in (Menneteau et al., 2019). The Thermofisher OrbiTrap Fusion was used as in (Payros et al., 2021).

### Mass spectrometry data analysis

The raw files generated during this work and previous studies (Wan et al., 2015; Wessels et al., 2016; Müller et al., 2020) were analyzed with MaxQuant (Cox et al., 2014) version 1.6.15.0. The minimal peptide length was set to 7. Trypsin/P, GluC or Chymotrypsin were used as the digestive enzymes. Search criteria included carbamidomethylation of cysteine as a fixed modification, oxidation of methionine and N-terminal acetylation as variable modifications. Up to two missed cleavages were allowed. The mass tolerance for the precursor was set to 20 ppm and 4.5 ppm for the first and the main searches, respectively, and 20 ppm for the fragment ions for Thermofisher instruments. The mass tolerance for the precursor was 0.07 and 0.006 Da for the first and the main searches, respectively, and for the fragment ions was 50 ppm and TOF recalibration was enabled for the Sciex TripleTOF 6600 instrument. The raw files were searched against the OpenProt fasta *Drosophila melanogaster* database (release 1.6, Altprots, Isoforms and Refprots). For the identification of RefProts, default MaxQuant settings were used (1% FDR both at the protein and PSM levels). Regarding AltProts identification, a minimum score of 70 was set for both modified and unmodified peptides (corresponding to the first quartile of the distribution of the score of RefProts from an analysis of the raw files with MaxQuant default settings). Candidates were filtered to obtain an FDR of 1% at the peptide level. Because alternative proteins are generally shorter than canonical proteins no FDR was set at the protein level and no filter was applied to the number of peptides per protein. A minimum sequence coverage of 70 % of the peptide sequence was required for alternative proteins identification. MSMS spectra were manually inspected by two independent operators. Peptides matching both a novel predicted protein and a RefProt were discarded. As implemented in OpenProt (Brunet et al., 2019), peptide matching two AltProts, two novel Isoforms or an AltProt and a novel Isoform were assigned to both proteins in each case. For quantification, the match between runs and iBAQ modules of MaxQuant were enabled. Quantitative comparison between AltProts and RefProts were performed on samples from the high pH reverse phase experiments only (Protein extraction and alternative protein enrichment protocol number 2, and data from (Müller et al., 2020)). As iBAQ represents an approximation of the absolute abundance of a protein (Fabre et al., 2014) and given the low number of observable peptides for AltProts, we considered that an AltProt was more (or less) abundant than its corresponding RefProt if the ratio between their iBAQ values was at least 10-fold different. Otherwise, AltProt and RefProt were considered to have similar expression levels. STRING v11.5 (Szklarczyk et al., 2021) was used for network generation and GO term / KEGG pathway analysis.

### Data availability

All the mass spectrometry data have been deposited with the MassIVE repository with the dataset identifier: MSV000088656.

### Confocal microscopy

For imaging experiments, S2 cells were co-transfected using actin-GAL4 driver with UAS-CG34150-GFP and UAS-AltProtCG34150-RFP or UAS-CG2650-GFP and UAS-AltProtCG2650-RFP (both constructions encoding AltProts also contain the start codons and sequence of the canonical proteins). Cells were transfected with effectene (Qiagen) according to manufacturer specification and as described in (Montigny et al., 2021). 48h post transfection, cells were fixed in 4% formaldehyde in phosphate buffer saline (PBS) at room temperature for 30 minutes. They were rinsed 3 times in PBS for 10 minutes. Nuclei were stained with DAPI and samples were rinsed several times in PBS. Coverslides were mounted in Prolong (Invitrogen) and images were acquired with a SP8 Leica confocal microscope. Three biological replicates were performed for each condition.

### Bioinformatic analyses

#### Detection of the AltProt and RefProt domains

The RefProt domains have been collected from the InterPro database (Blum et al., 2021a). All domains identified on UniProtKB reviewed proteins of *Drosophila melanogaster* (Proteome identifier: UP000000803) have been recovered using the EBI REST API.

The domains on AltProt sequences have been identified using InterProScan (Jones et al., 2014)(v5.52-87.0) looking for signatures in the Pfam database (Mistry et al., 2021). The signatures with an e-value lower than 10^−5^ have been selected. Pfam identifiers have been mapped to InterPro accessions using the InterPro cross-references collected through the EBI REST API.

#### Detection of the short linear motifs (SLiMs)

Classes of SLiMs have been downloaded from the Eukaryotic Linear Motif (ELM) database (Kumar et al., 2020). Classes with a pattern probability lower than 0.01 and having at least one true positive instance detected in D. melanogaster in ELM database have been selected.

The short linear motifs (SLiMs) have then been detected in the disordered regions of the AltProts using the IUPred2A (Mészáros et al., 2018a) and the Short Linear Motif Probability tool (SLiMProb) of SLiMSuite (Edwards et al., 2020), using the following SLiMProb parameters: iumethod = long, iucut = 0.2, minregion = 5.

#### Associations between SLiM and domain usage and AltProts classes

To check whether AltProt classes were preferentially associated with SLiM or domain usage, Chi-squared tests of independence have been performed.

#### SLiM enrichments and depletions among AltProt classes

For each class type of motif (LIG, DOC, TRG, MOD, CLV, DEG), and for each class of AltProt (ncRNA, Isoform, 5'UTR, CDS, 3'UTR), enrichment and depletion in AltProt with at least one motif of the class type among the AltProt of the class have been assessed, using one-sided Fisher's exact tests. The p-values computed have then been adjusted for multiple comparisons using a Benjamini-Hochberg procedure.

For each class of motif, and for each class of AltProt (ncRNA, Isoform, 5'UTR, CDS, 3'UTR), enrichment and depletion in AltProt with at least one motif of the class among the AltProt of the class have been assessed, using one-sided Fisher's exact tests. The p-values computed have then been adjusted for multiple comparisons using a Benjamini-Hochberg procedure.

Disorder regions (sequence of at least 5 amino acids) were predicted using IUPred2A (Mészáros et al., 2018b) using the long disorder setting. Prediction of transmembrane helices and signal peptides were performed using TMHMM - 2.0 (Krogh et al., 2001) and SignalP - 5.0 (Almagro Armenteros et al., 2019), respectively. DeepLoc (Almagro Armenteros et al., 2017) was used to predict AltProts subcellular localization.

## Results and discussion

### Genome wide identification of Alternative proteins in Drosophila melanogaster

In order to identify new alternative proteins in *Drosophila melanogaster* we developed a customized peptidomics workflow (Figure 1). We used a combination of protein extraction and small proteins enrichment protocol as it was previously shown to increase the number of AltProts identified by mass spectrometry (Ma et al., 2016; Cardon et al., 2020). Extensive fractionation, using high pH reverse phase chromatography, as well as specific enrichment of short proteins, through SDS-PAGE, ultrafiltration, acid precipitation and reverse phase chromatography, was employed to retrieve AltProts from adult flies, 0-24h embryos and S2 cells (Figure 1). We also re-analyzed MS data available in public repositories. In total, 1,068 MS files were analyzed using optimized MaxQuant parameters and the OpenProt predicted AltProts database (Figure 1). In total, 401 AltProts as well as 8,615 RefProts (including 267 RefProts containing less than 100 amino acids annotated in UniProtKB) were identified (Figure 2 and Supplementary table 1 and 2). The identification scores obtained for the AltProts were similar to the ones measured for a typical proteomics analysis (median Andromeda score of 98.52 for AltProts and 101.97 for RefProts) (Supplementary Figure 1A). The majority of the AltProts identified here are short proteins (88.8% of AltProts are less than 150 amino acids) (Supplementary Figure 1B). Comparing the AltProts identified to the ones with MS evidence in OpenProt and a recent article (Wang et al., 2022), only 2 were common to the three datasets, 29 were found in at least two datasets and 374 new AltProts were identified in this study (Supplementary Figure 1C). The low overlap between the datasets might be explained by the different sample types and extraction and fractionation protocols used (Cardon et al., 2020). Amongst the 401 non-annotated proteins identified, 30 were new Isoforms (Figure 2). As defined in OpenProt (https://www.openprot.org/), we refer here as Isoform (or novel Isoform) any non-annotated proteins that share some homology with a RefProt (either partially overlapping coding sequences, although only Isoform unique peptides are used for their identification). Next, we looked at the RNA types and regions from which AltProts are produced (Figure 2). We used a recently suggested nomenclature (Mudge et al., 2021) to refer to the types of ORFs encoding the AltProts (Figure 2). Surprisingly, whereas pioneering studies identified non-coding RNA or untranslated regions of mRNAs as main sources of AltProts (Plaza et al., 2017b), the majority of AltProts identified in our study are produced from mRNA and more particularly from alternative reading frames in canonical CDS (intORFs). With more than 300 AltProts produced from uORFs, intORFs or dORFs, our data strongly advocate towards a polycistronic nature of mRNA in *Drosophila melanogaster* (Figure 2). Of note, 52 AltProts are produced from previously predicted long non-coding RNA (Figure 2), including 1 AltProt encoded by a precursor of miRNA (pri-miRNA) (Figure 2), supporting the idea that miPEPs (miRNA-encoded peptides) are expressed in flies (Immarigeon et al., 2021; Montigny et al., 2021). Regarding the sources of production of AltProts, and more particularly the chromosomes they are produced from, a distribution similar to the predicted AltProts distribution from OpenProt was observed (Supplementary Figure 2A-B), although slight differences could be noticed. The proportion of AltProts produced from the chromosomes 2R and 3L were higher than expected contrary to the chromosomes 4 and X where a lower proportion of AltProts were identified (Supplementary Figure 2A-B). Interestingly, the proportion of new Isoforms as well as Altprots synthesized from uORFs and lncRNA-ORFs were more represented than expected (Supplementary Figure 2C-H).

**Figure 1:**
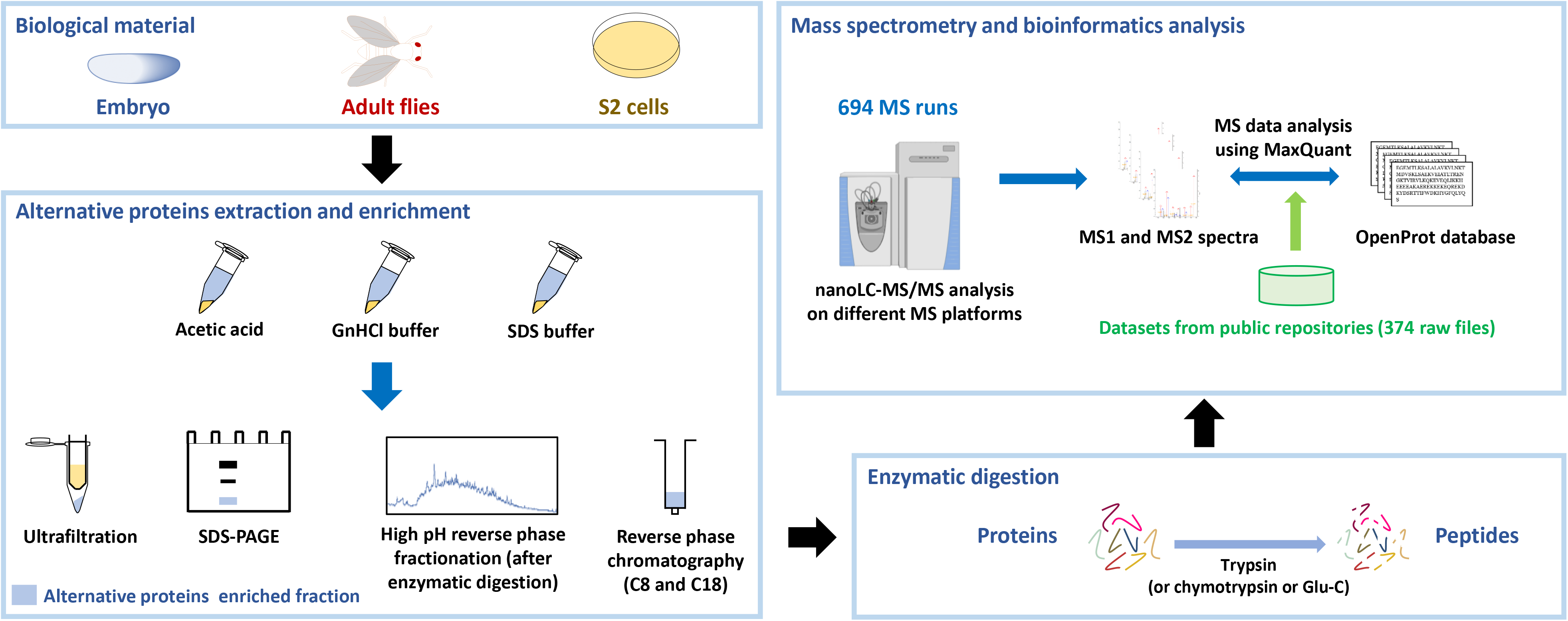
Peptidomics workflow to identify alternative proteins in *Drosophila melanogaster*. Proteins were extracted from embryos, adult flies or S2 cultured cells using different extraction protocols. Alternative proteins were then enriched from the total protein pool and digested with trypsin (or other enzymes). The resulting peptides were injected on different mass spectrometry platforms and the generated MS data, as well as datasets available from public repositories, were analyzed using MaxQuant with the OpenProt database.

**Figure 2:**
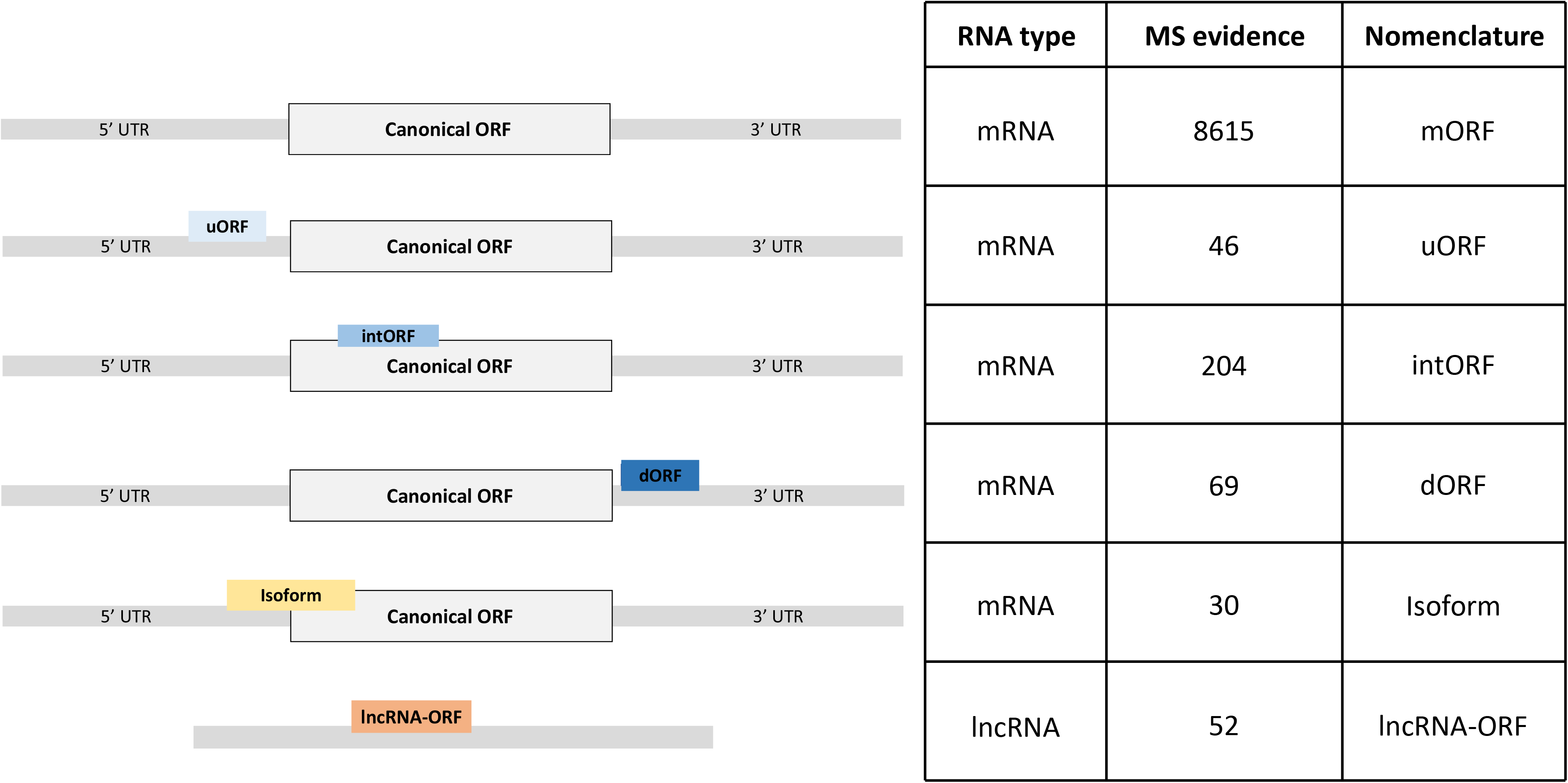
Different classes of ORFs produce yet non-annotated alternative proteins. Hundreds of alternative proteins are produced from ORFs from novel isoforms, ORFs located in the 5’UTR (uORF), coding sequence (intORF) or 3’UTR (dORF) of mRNAs, or from non-annotated ORFs on long non-coding RNAs (lncRNA-ORF). The nomenclature of the different classes of ORFs was adapted from (Mudge et al., 2021).

We next looked at the position of the start codon of the 401 AltProts identified. Surprisingly, only 16.7% of the AltProts identified here are produced from the first predicted start codon (Figure 3A). As expected AltProts produced from uORFs are synthesized from the first start codon more frequently than AltProts produced from intORFs and dORFs (45.7% versus 5.9% and 0%, respectively) (Figure 3B-D and Supplementary Figure 3). Interestingly, new Isoforms and AltProts produced from lncRNA-ORFs follow a pattern similar to uORFs with 40% and 42.3% of these proteins being synthesized from the first start codon (Figure 3B, 3E-F and Supplementary Figure 3). These data highlight that, although the translation of AltProts from the first ORF on an RNA is the most probable (notably for AltProts produced from lncRNA-ORFs), 334 of the newly proteins identified here are translated from further ORFs on RNAs. Notably, 22 AltProts are synthesized from the 20^th^ predicted ORF or beyond (Figure 3A).

**Figure 3:**
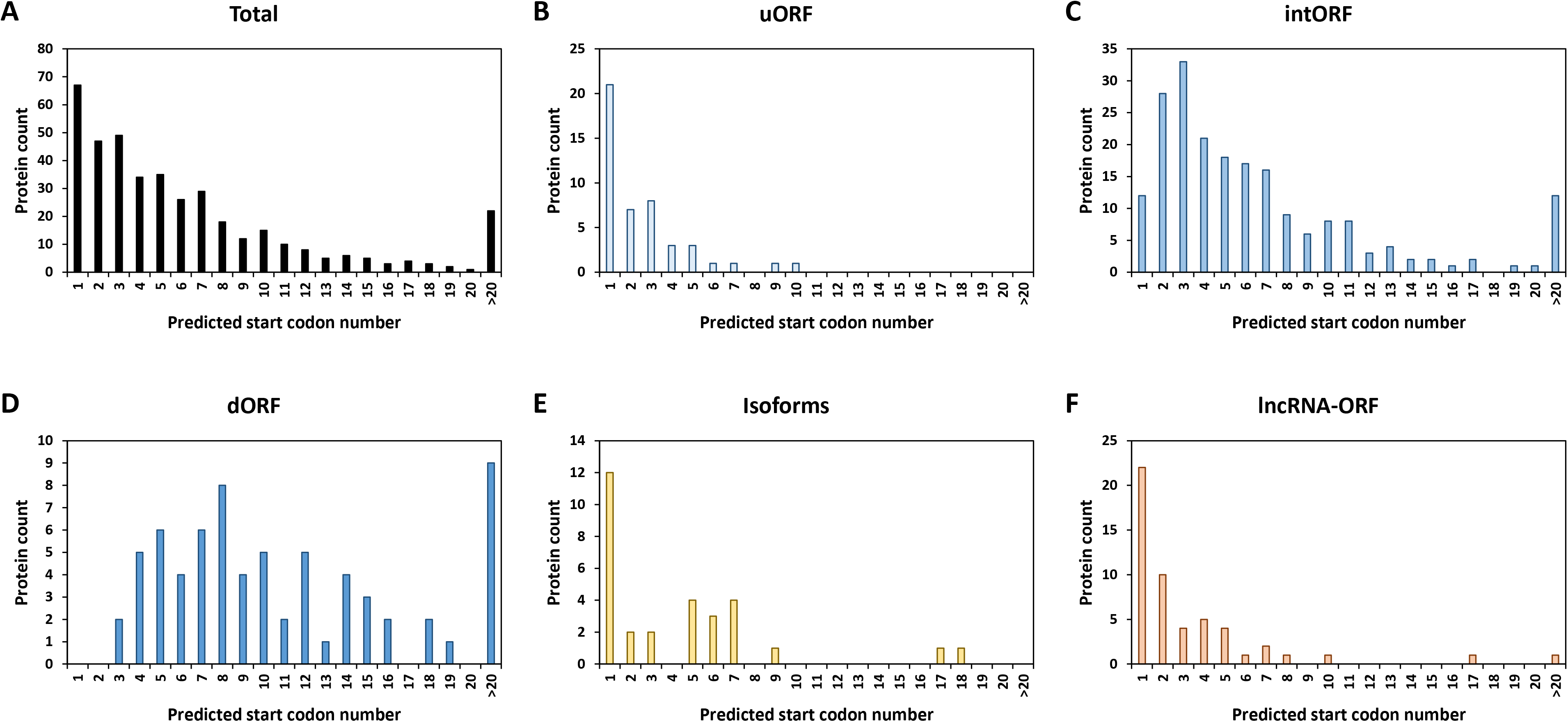
Alternative proteins are not mainly produced from the first predicted ORF. Distribution of the alternative proteins counts depending on the order of the position of the predicted start codons for all the newly identified proteins (Total) (A), isoforms (B), proteins identified from ORFs encoded by lncRNA (lnc-ORF) (C), uORF (D), AltCDS (E) or dORF (F) on mRNA.

### Structural properties of Alternative proteins in Drosophila melanogaster

Next, the chemical characteristics of the AltProts identified were investigated. First, we looked at the size distribution of the AltProts depending on the type of ORF they are synthesized from (Figure 4A). As expected Isoforms are longer than other AltProts (median length of 221.5 and 52 amino acids, respectively) (Figure 4A and Supplementary Figure 4A-B). Within AltProts, proteins produced from lncRNA-ORFs are slightly longer than the alternative proteins synthesized from mRNA (median length of 67, 49, 52.5 and 44 for AltProts from lncRNA-ORFs, uORFs, intORFs and dORFs, respectively) (Figure 4A and Supplementary Figure 4A-B).

**Figure 4:**
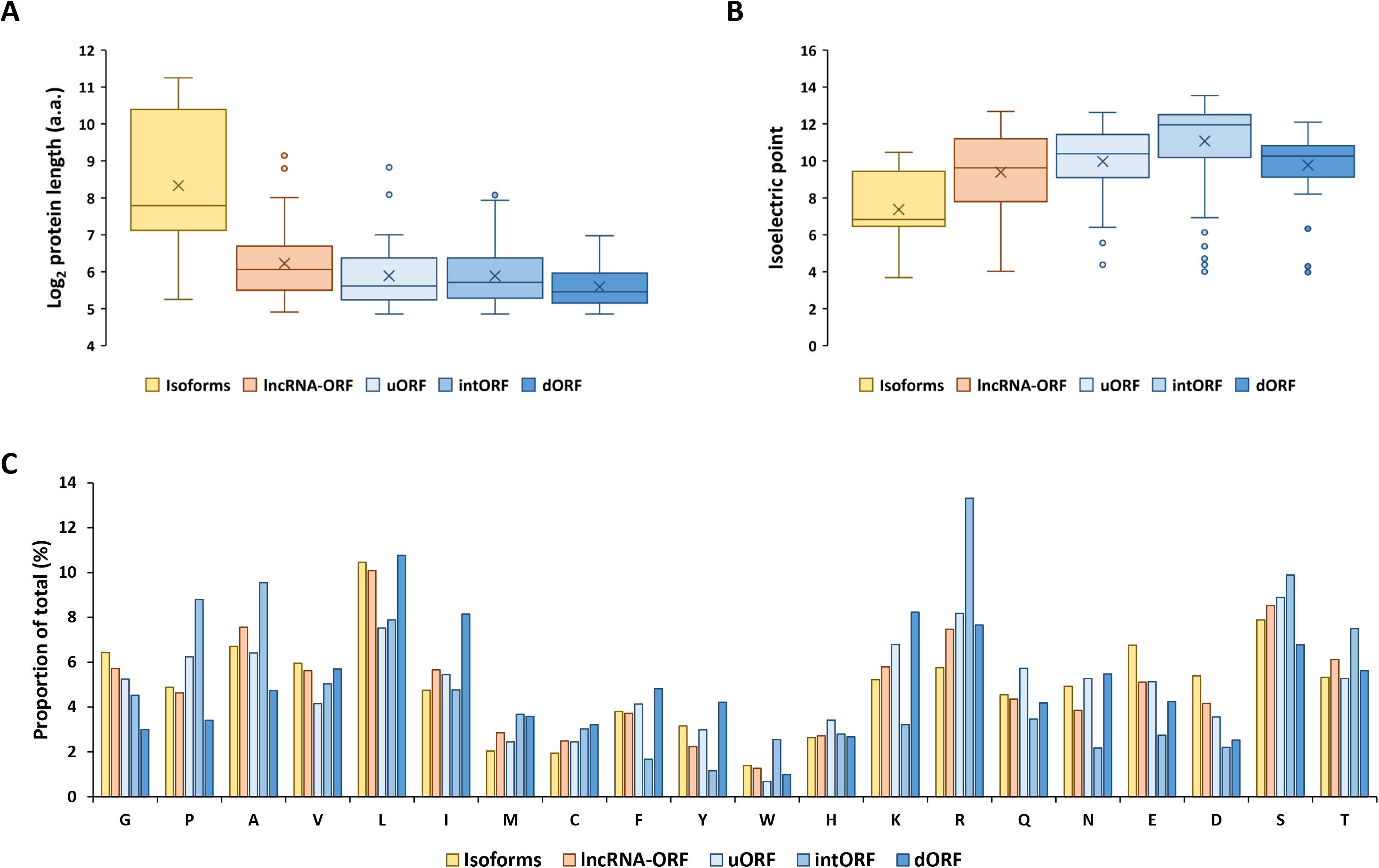
Alternative proteins produced from different classes of ORFs have different chemical properties. A. Distribution of the number of newly identified proteins in each type of ORF class and depending on their length in amino acids (a.a.). B. Repartition of the isoelectric point measured for the proteins identified in each type of ORF class. C. Amino acids proportions obtained from the sequences of the proteins identified in each type of ORF class.

Comparing the isoelectric point (pI) of the different class of AltProts revealed that Isoforms have lower pI than other AltProts (*p* < 9.04×10^−5^) (Figure 4B). In addition, AltProts produced from intORFs tend to have higher pI than the other AltProts (*p* < 0.0013) (Figure 4B). This might be explained by the fact that the overall amino acid composition of AltProts produced from intORFs differs from other AltProts (Figure 4C). The former has more arginine, alanine and tryptophan and less asparagine, lysine and glutamic acid (*p* < 2.2×10^−16^) (Figure 4C). This difference in composition might point toward specific functions of AltProts produced from intORFs.

We then performed a Gene Ontology (GO) analysis on the host genes, from which the AltProts are produced, to gain some insight in the possible functions of the newly discovered proteins (Figure 5A). Interestingly, the most significant term enriched were cell development (FDR = 2.4×10^−7^) and cell differentiation (FDR = 9.6×10^−7^), suggesting that the AltProts identified in this study might have functions related to developmental processes (Figure 5A and Supplementary Figure 5). These pathways are mainly enriched in host genes from AltProts produced from intORFs and dORFs (Supplementary Figure 6 and 7). No pathway was found enriched in AltProts produced from uORFs or Isoforms (Supplementary Figure 8).

**Figure 5:**
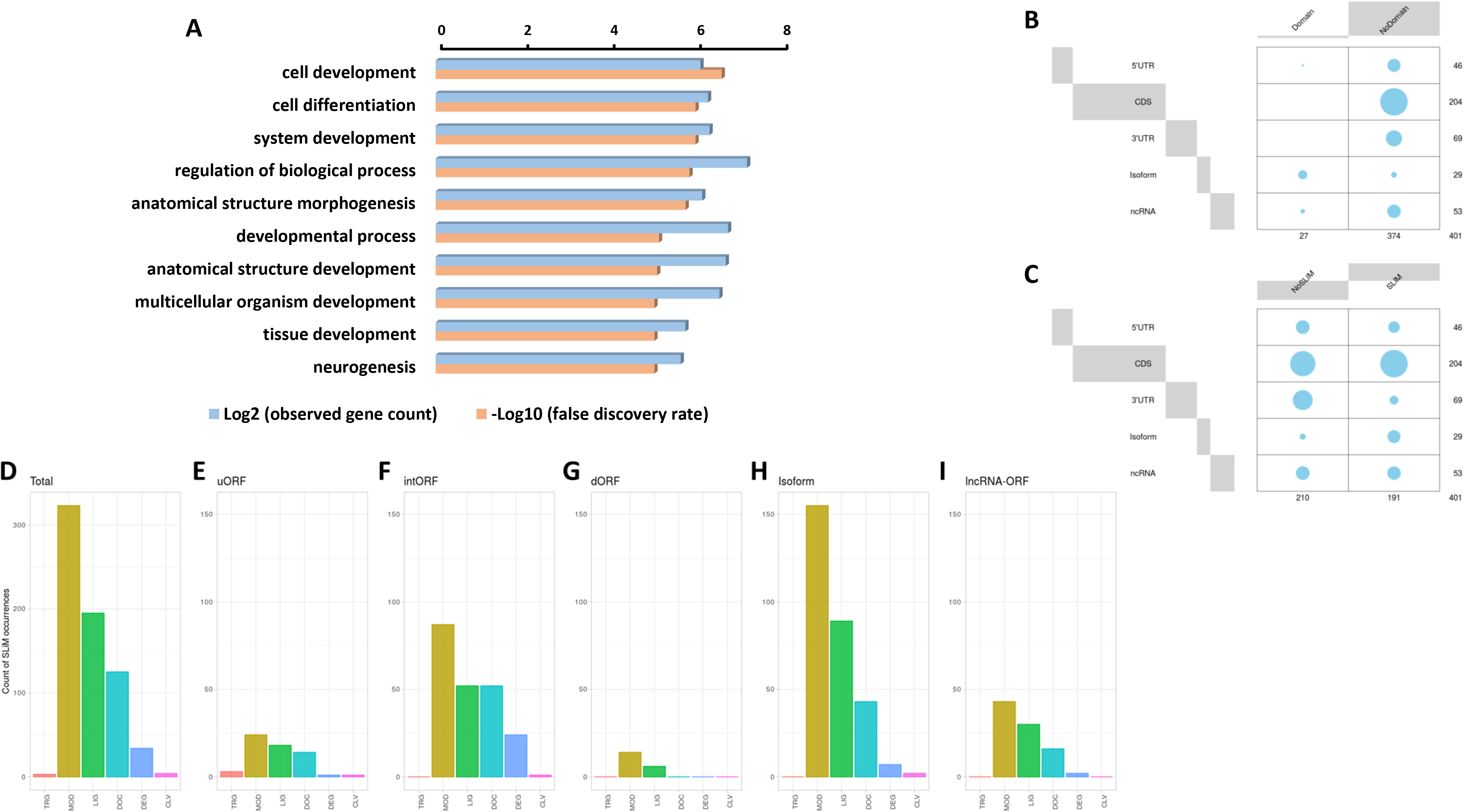
Identification of protein domains and short linear motifs in alternative proteins. A. Gene Ontology term analysis of the host genes of the alternative proteins identified. B. Balloon plots showing the presence or lack of protein domain for the different types of ORFs the AltProts are produced from. The area is proportional to the frequency. C. Balloon plots showing the presence or lack of SLiMs for the different types of ORFs the AltProts are produced from. The area is proportional to the frequency. D-I. Counts of the different classes of SLiMs in the AltProts identified in this study (D) or identified from uORFs (E), intORFs (F), dORFs (G), isoforms (H) or lncRNA-ORFs (I). The SLiMs classes are targeting sites for subcellular localization (TRG), post-translational modification sites (MOD), ligand binding sites (LIG), docking sites (DOC), degradation sites (DEG) and proteolytic cleavage sites (CLV).

In order to dig deeper into the possible role of the AltProts of *Drosophila melanogaster*, several prediction tools were used to identify potential protein domains, disordered regions or subcellular localization signals. Looking at protein domains, InterPro (Blum et al., 2021b) predicted that 27 of the AltProts identified might have one or more protein domain (Figure 5B and Supplementary table 3). Analysis using the TMHMM - 2.0 (Krogh et al., 2001) and SignalP - 5.0 (Almagro Armenteros et al., 2019) software identified possible transmembrane domains and signal peptides for 33 and 9 AltProts, respectively. Interestingly, the AltProt IP_1410397 encoded by the host gene CG15784 had both a signal peptide and a transmembrane domain predicted (both with probabilities > 0.8) in the first 30 amino acids of its sequence (Supplementary Figure 9).

We next looked for the presence of short linear motifs (SLiMs) in the AltProts identified. SLiMs are functional short stretches of protein sequence that are generally involved in protein-protein interactions (Hraber et al., 2020). A total of 684 SLiMs were mapped on 191 AltProts (Figure 5C-D and Supplementary table 4). Most of the SLiMs retrieved belong to the post-translational modification sites (MOD) (enriched in isoforms, Benjamini-Hochberg adjusted p-value = 3.5 .10^−4^, odd ratio = 5.41), ligand binding sites (LIG) (especially enriched in Isoforms, Benjamini-Hochberg adjusted p-value = 1.42 .10^−7^, odds ratio = 11.04) and docking sites (DOC) classes (47%, 29% and 18% of the SLiMs identified, respectively) (Figure 5D, Supplementary Figure 10A and Supplementary table 4). The SLiMs the most represented were Polo-like kinase1 and 4 phosphosites motifs (MOD_PlK_1 and MOD_Plk_4, found on 64 and 100 AltProts, respectively), cyclin N-terminal domain docking motifs (DOC_CYCLIN_RXL_1, found on 38 AltProts) and Atg8 protein family ligands motifs (LIG_LIR_Gen_1, found on 51 AltProts) (Supplementary Figure 10A and Supplementary table 4), suggesting a possible role of these AltProts in drosophila cell cycle and autophagy. Importantly, Isoforms were the only class of AltProts which displayed a significant enrichment in SLiMs (Supplementary Figure 10B-C and Supplementary table 5). Looking at each type of ORFs, slight differences in SLiMs classes could be observed (Figure 5E-I). The SLiMs class targeting sites for subcellular localization (TRG) was identified only on 1 AltProt produced from an uORF (Figure 5E). It was surprising to notice that AltProts from dORFs only have 20 SLiMs detected on 11 AltProts (Figure 5G). In addition, this type of AltProts do not seem to bear any protein domain and although all the SLiMs identified are from the MOD and LIG classes, the number of SLiMs from these classes detected is still lower than expected (Benjamini-Hochberg adjusted p-value = 0.009 and 0.0006, odds ratio = 0.35 and 0.17, respectively) (Supplementary table 5). These results suggest that most of the AltProts produced from dORFs are less susceptible than other AltProts to carry particular functions based on domain prediction and that the ORF itself might be mainly involved in the regulation of protein translation as recently proposed (Wu et al., 2020).

Next, the IUPred2A software was used to predict disordered regions within AltProts. Around 50% of the AltProts identified in our study contained one or more predicted disordered region (Supplementary Figure 11). A higher proportion of Isoforms (70%) and AltProts produced from uORFs (58.7%) and intORFs (55.9%) tend to have disordered regions compared to lncRNA-ORFs (38.5%) or dORFs (26.1%) (Supplementary Figure 10). This is in agreement with a recent report in plants, which also predicted that numerous non-annotated short proteins might contain disordered regions, transmembrane domains or signal peptides (Fesenko et al., 2021).

DeepLoc (Almagro Armenteros et al., 2017) was used to predict possible subcellular localization of Altprots. A potential localization was assigned to 79 out of 401 with a probability higher than 0.8 (Figure 6A and Supplementary table 5). Surprisingly, 35 of these AltProts were predicted to be mitochondrial, 18 extracellular and 17 nuclear (Figure 6B and Supplementary table 6). Only 7 Altprots are predicted to be cytoplasmic, 1 potentially localized in the Golgi and 1 in the endoplasmic reticulum (Figure 6B and Supplementary table 6). When comparing the predicted subcellular localization of Altprots produced from mRNAs and their corresponding RefProts, only 18% (12 out of 67) were concordant (Supplementary Figure 12). In order to validate the prediction from DeepLoc, the AltProt and RefProt of the gene *CG34150*, tagged with a red fluorescent protein (RFP) and a green fluorescent protein (GFP) respectively, were transfected in S2 cells and co-expressed under the same *actin* promoter. Confocal imaging revealed that, in agreement with the DeepLoc prediction, both proteins are colocalized in S2 cells (Figure 6C). Similarly tagged versions of the AltProt and RefProt of the gene *CG2650*, for which no subcellular localizations were predicted in animal cells, were also expressed in S2 cells (Figure 6D). Surprisingly, colocalization could be observed in certain cells whereas other cells showed different localization patterns between the two proteins in the S2 cells within the same experiment (Figure 6D). This might be indicative that the AltProt and RefProt of CG2650 are colocalized upon particular cellular conditions (e.g. specific cell cycle stages…). These experiments also showed that the two *CG34150* and *CG2650* AltProts are expressed despite the presence of the ATG of the canonical ORF, confirming peptide detection observed in MS analysis.

**Figure 6:**
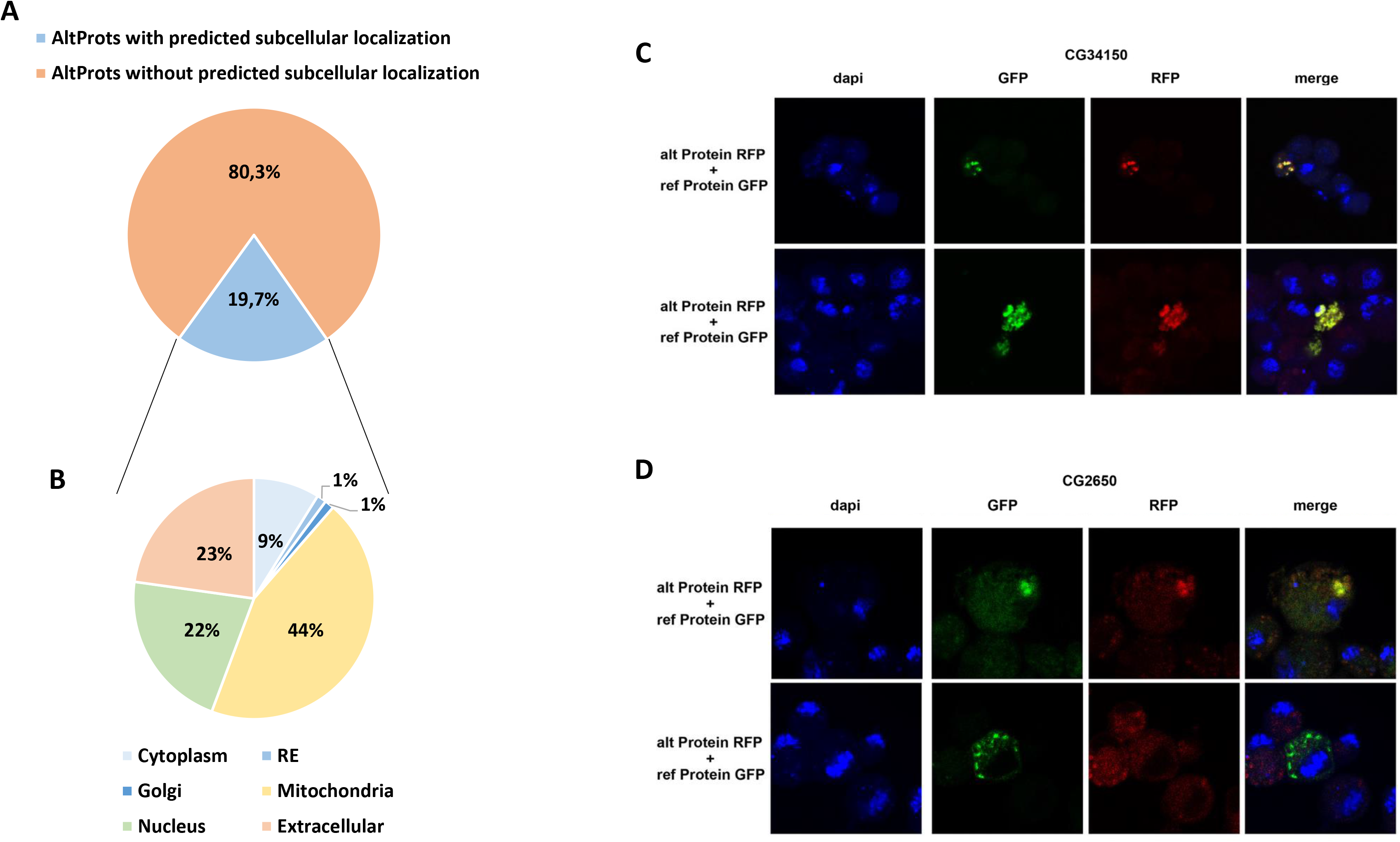
Prediction of alternative proteins subcellular localization. A. Proportion of AltProts with or without predicted subcellular localization. B. Proportion of AltProts predicted to be localized in different organelles. C. Confocal microscopy analysis of the co-expression of the AltProt (alt Protein) (with the sequence of the RefProt before the AltProt starting codon) and RefProt (ref Protein) encoded by the CG34150 gene, tagged with RFP and GFP, respectively, in S2 cells (stained with DAPI). D. Confocal microscopy analysis of the co-expression of the AltProt (alt Protein) (with the sequence of the RefProt before the AltProt starting codon) and RefProt (ref Protein) encoded by the CG2650 gene, tagged with RFP and GFP, respectively, in S2 cells (stained with DAPI).

Overall, these data corroborate previous observations in human suggesting that AltProts might have independent functions or roles related to their corresponding RefProts (Chen et al., 2020a). Here we identified 235 AltProts for which at least a protein domain, a SLiM or a subcellular localization was predicted (Figure 5B and 5D and Figure 6A). Although further functional experiments would be necessary to better understand the role of these AltProts, these predictions provide first hints regarding the functions of the Altprots identified in this study.

### Alternative proteins are not necessarily less abundant than canonical proteins

We next wondered if AltProts can be more abundant than their corresponding RefProts as previously shown for the human alternative protein altMiD51 (Delcourt et al., 2018). Comparing the intensities measured for peptides from AltProts and RefProts from total lysates and high pH reverse phase fractionation (no specific AltProts enrichment, see material and methods section protocol 2 as well as data from (Müller et al., 2020)), the peptides from the latter were slightly more intense (1.96 fold difference of the average peptides intensities measured for AltProts and RefProts, Supplementary figure 13A). This implies that some AltProts might be as abundant as RefProts in drosophila. The iBAQ values, which represent an approximation of the abundance of a protein (Krey et al., 2014), were used to compare the abundance of AltProts with the abundance of their corresponding RefProts in total lysates and high pH reverse phase fractionation experiments (Figure 7A and 7B). Out of the 39 pairs of AltProts/RefProts for which iBAQ values were measured, 22 did not show any difference in abundance between AltProts and RefProts expression levels (less than 10-fold difference between the iBAQ values), whereas 11 AltProts were more abundant (Figure 7A and 7B). Only 6 RefProts were more abundant than their corresponding AltProts (Figure 7A and 7B). This trend was observed in two independent datasets (Supplementary figure 13B) and we did not observe any bias based on the length of the AltProts (Supplementary figure 14). These data reveal that, in several cases, alternative proteins are actually the main protein produced from their corresponding genes.

**Figure 7:**
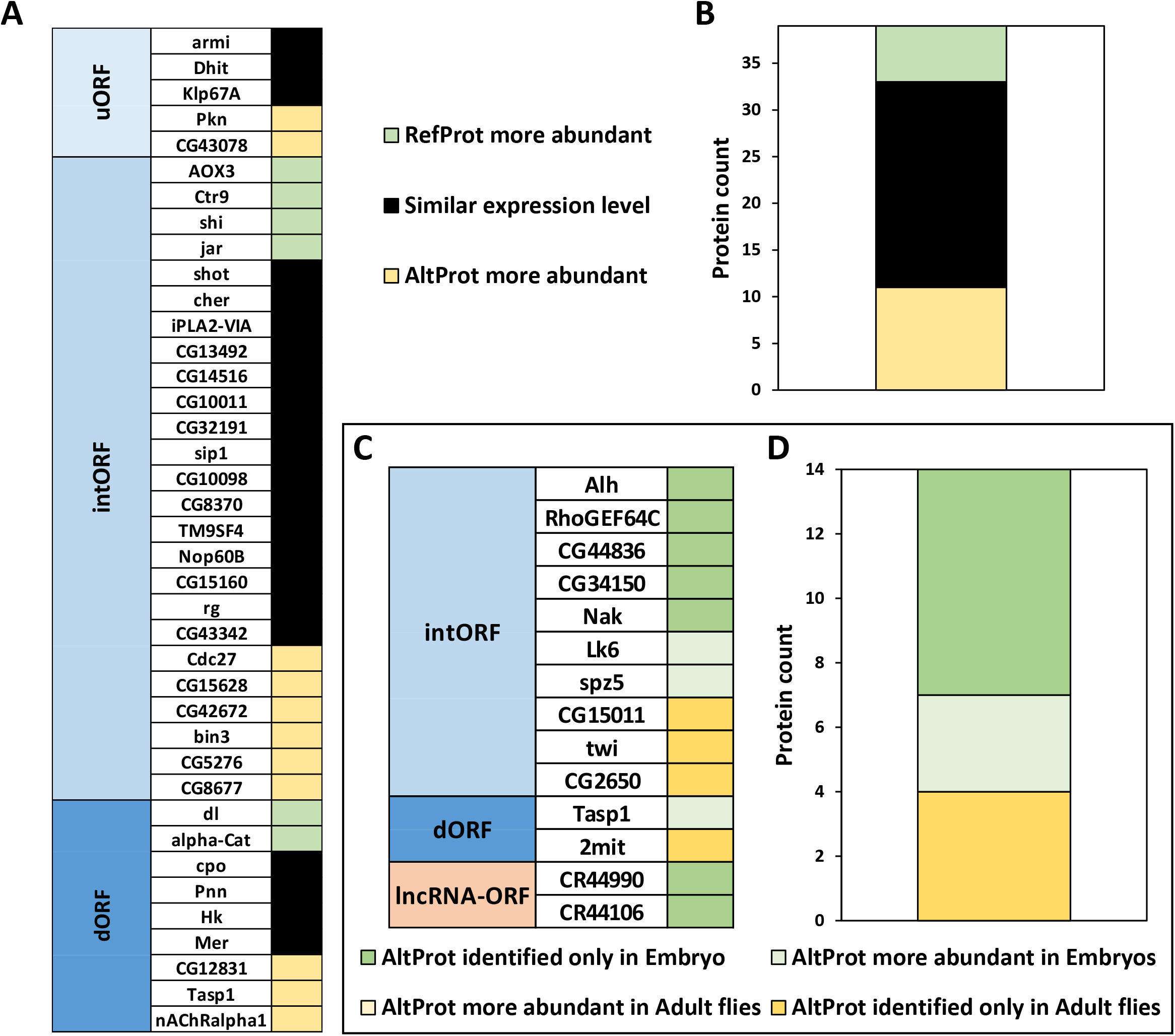
Alternative proteins can be more abundant than their corresponding canonical proteins and their expression is developmentally regulated. A. Comparison of the approximation of the relative abundance measured for each couple of canonical (RefProts) and alternative (AltProts) proteins for each type of ORF using the intensity based absolute quantification approach (iBAQ). An AltProt/RefProt is considered more abundant than its counterpart if its iBAQ value is at least 10-fold higher. B. Graph showing the protein counts of AltProts more abundant than RefProts, RefProts more abundant than AltProts and similar expression levels between AltProts and RefProts. C. The fold change between embryo and adult fly was calculated for each AltProt and normalized by the maximum intensity between the two samples. An AltProt/RefProt is considered more abundant in a specific developmental stage if its intensity value is at least 3-fold higher. D. Graph showing the protein counts of AltProts more abundant, or identified only, in embryo or in adult flies.

### Developmentally timed and stress specific production of alternative proteins

Next, the expression of AltProts was compared to monitor potential changes between embryos and adult flies (material and methods protocol 1 and 2, Figure 7C-D). All the 14 AltProts for which we obtained quantitative data in at least two biological replicates were more abundant in one developmental stage (Figure 7C-D). Three AltProts were identified both in embryos and adult flies but were at least 3 times more abundant in embryos (Figure 7C-D). The remaining 11 AltProts were identified only in one stage (Figure 7C-D), suggesting that the expression of most of the AltProts quantified here are developmentally timed. Four AltProts were identified only in adult flies whereas seven were specific to embryo samples, including two AltProts produced from lncRNA-ORFs (Figure 7C-D).

We also tested whether the expression of AltProts varies upon stress. Embryos were treated with heat-shock at 37°C for up to 3 hours or kept at 25°C and analyzed to identify alternative proteins. We were able to identify 22 AltProts in these samples, including 10 AltProts that were identified only in heat-shock treated embryos (Supplementary table 7). These results demonstrate that alternative proteins are produced under specific developmental stages or stress conditions in *D. melanogaster*.

## Conclusion

Recent studies in human suggested that the complexity of the genome was underestimated (Brunet et al., 2021; Ouspenskaia et al., 2021) and that many unannotated proteins might fulfill important functions, related or not to canonical proteins (Plaza et al., 2017b; Chen et al., 2020a). However, it is not clear whether this is specific to human or whether this characteristic is present in every species since we still lack deep analysis of this alternative proteome in many species, including the model organism *D. melanogaster*. In flies, until now, mainly data from ribosome profiling experiments were available to annotate putative translated alternative sORF (Aspden et al., 2014; Patraquim et al., 2020). In the present study we developed a deep peptidomics workflow which combines several extraction methods and enrichment protocols with mass spectrometry and dedicated bioinformatics analysis to identify new alternative proteins in flies. We proved for the first time the existence of 374 AltProts predicted in OpenProt (Figure 2), significantly increasing the repertoire of not yet annotated proteins in *D. melanogaster*. Many of these AltProts even escaped from ribosome profiling experiments as they are encoded by alternative frames within the annotated CDS. Contrasting with these results, we did not find many unannotated proteins with a coding sequence of more than 100 codons, revealing that the annotation of proteins with ORF of 100 codons or more is precise and reliable. On the other hand, our study shows that many proteins of less than 100 amino acids remain to be discovered, especially considering the fact that we did not search for alternative proteins of less than 30 amino acids, which are known to be expressed and functional in *D. melanogaster* (Magny et al., 2013; Zanet et al., 2015; Immarigeon et al., 2021; Montigny et al., 2021) and would require further investigation. Interestingly, these AltProts are not necessarily produced from the first predicted ORF on an RNA (Figure 3), one spectacular result came from an AltProt being synthesized from the 134^th^ predicted ORF on the dumpy mRNA (Supplementary table 1). Another key observation is that more than 300 mRNAs are actually polycistronic (Figure 2). The main source of production of AltProts in *Drosophila melanogaster* is alternative frames in canonical coding sequences (intORFs) (Figure 2) possibly a specificity of drosophila since in human and mouse, AltProts are produced mainly from lncRNA (https://www.openprot.org/). Surprisingly, AltProts produced from intORFs have different amino acids composition compared to other AltProts, pointing toward a possible specific function of this type of AltProts (Figure 4B-C). Regarding potential functions of the identified AltProts, protein domain, SLiMs or subcellular localization were predicted for 235 of them (Figure 5B-C and 6A) pointing toward potential functions for these small proteins. However, the lack of predicted protein domains and low number of SLiMs identified on dORFs implies that the AltProts produced from these ORFs might not be functional. Fluorescence confocal microscopy confirmed the colocalization of the *CG34150* AltProt and RefProt and showed that the *CG2650* AltProt and RefProt can colocalize upon certain conditions (Figure 6C-D). The comparison of the abundance (using the iBAQ value as an approximation) of alternative and canonical proteins revealed that AltProts are not necessarily less abundant and might actually be the main product of several genes (Figure 7A and 7B). These data suggest that it might be worth reconsidering the phenotypes observed in certain mutants in *D. melanogaster* as they might be mediated by the mutation/deletion of the alternative protein rather than the canonical one. Finally, several AltProts were identified in only specific developmental stages or upon heat shock, implying that their expression is finely tuned during D. melanogaster development or under stress conditions (Figure 7C and 7D). These proteins might have important functions during development or heat shock response, hence requiring further functional investigation.

## Acknowledgements

This work has been supported by the Fondation ARC pour la recherche sur le cancer. B.F. is funded by Biotechnology and Biological Science Research Council (ref: BB/L002817/1) and a long term EMBO fellow (ALTF 1204-2015) cofounded by Marie Curie Actions (LTFCOFUND2013, GA-2013-609409). S.A.C. is funded by a Fondation pour la Recherche Médicale fellowship (FDT202106013072). D.K. is funded by Biotechnology and Biological Science Research Council (ref: BB/L002817/1). The work was funded in part by grants from the Région Occitanie, European funds (Fonds Européens de Développement Régional, FEDER), Toulouse Métropole, and the French Ministry of Research with the Investissement d’Avenir Infrastructures Nationales en Biologie et Santé program (ProFI, Proteomics French Infrastructure project, ANR-10-INBS-08). We acknowledge the Centre de Calcul Intensif d’Aix-Marseille for granting access to its high-performance computing resources.

## Conflicts of Interest

The authors declare that the research was conducted in the absence of any commercial or financial relationships that could be construed as a potential conflict of interest.

## Notes

### Competing Interest Statement

The authors have declared no competing interest.

