## Supplementary_data for "In depth exploration of the alternative proteome of *Drosophila melanogaster*"

Supp Figure 1

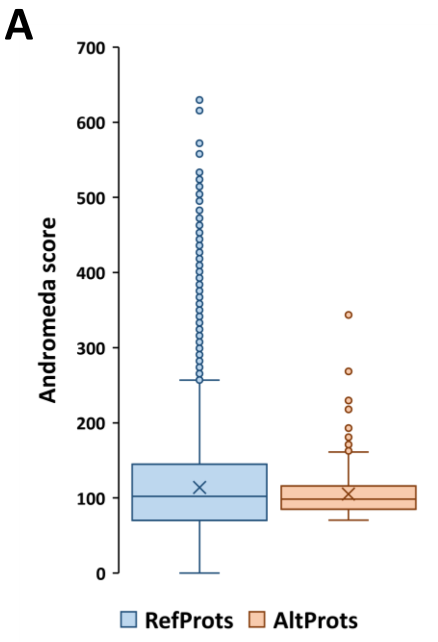


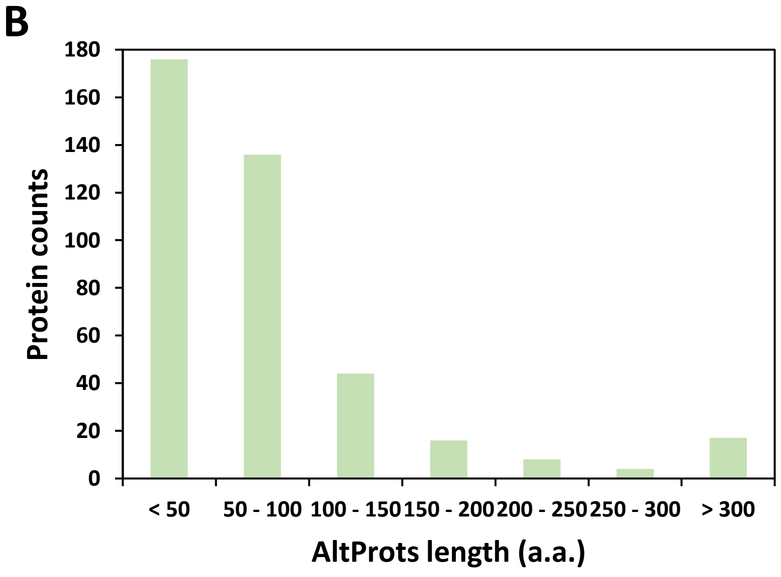


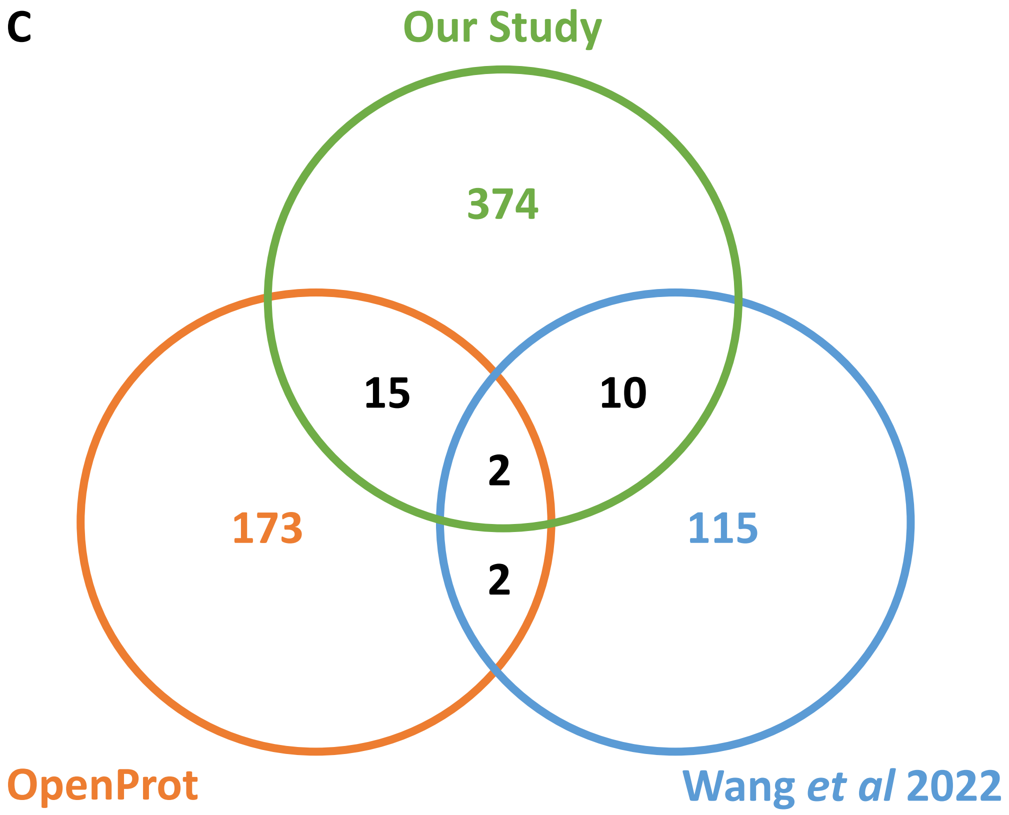

Supp Figure 2

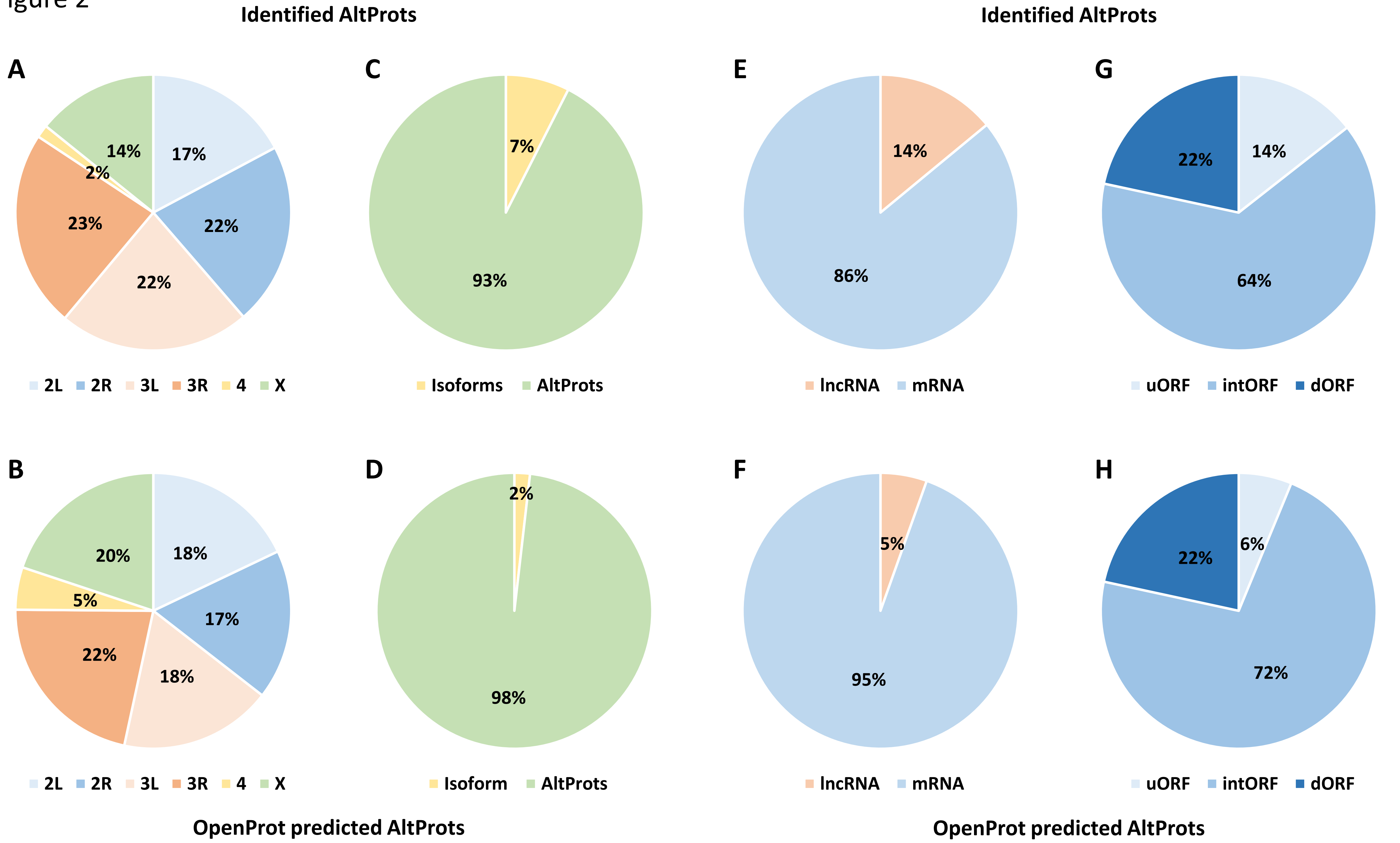

Supp Figure 3

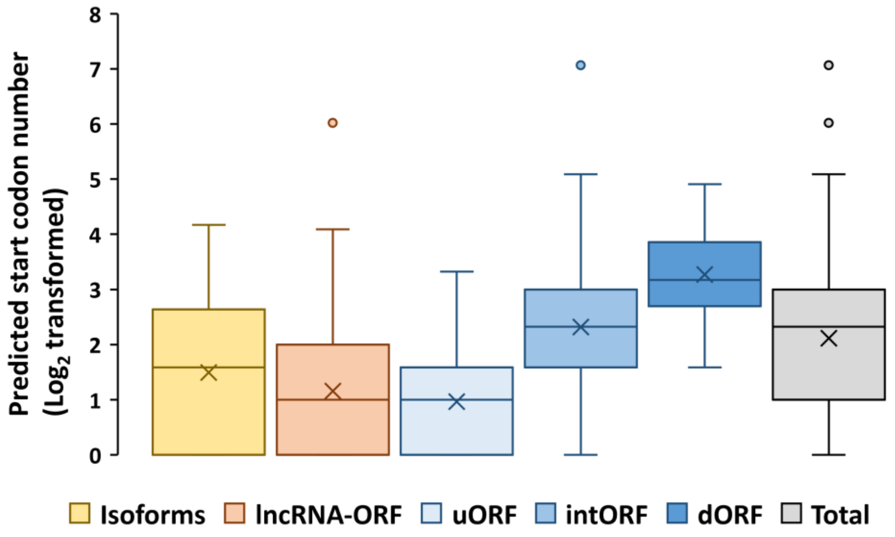

Supp Figure 4

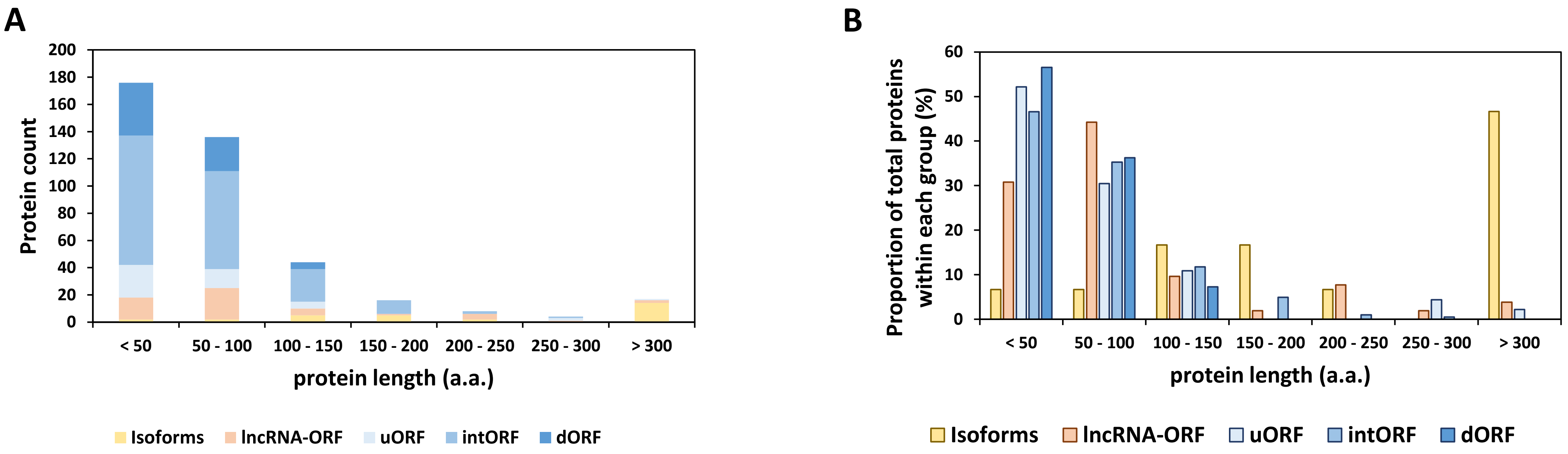

Supp Figure 5

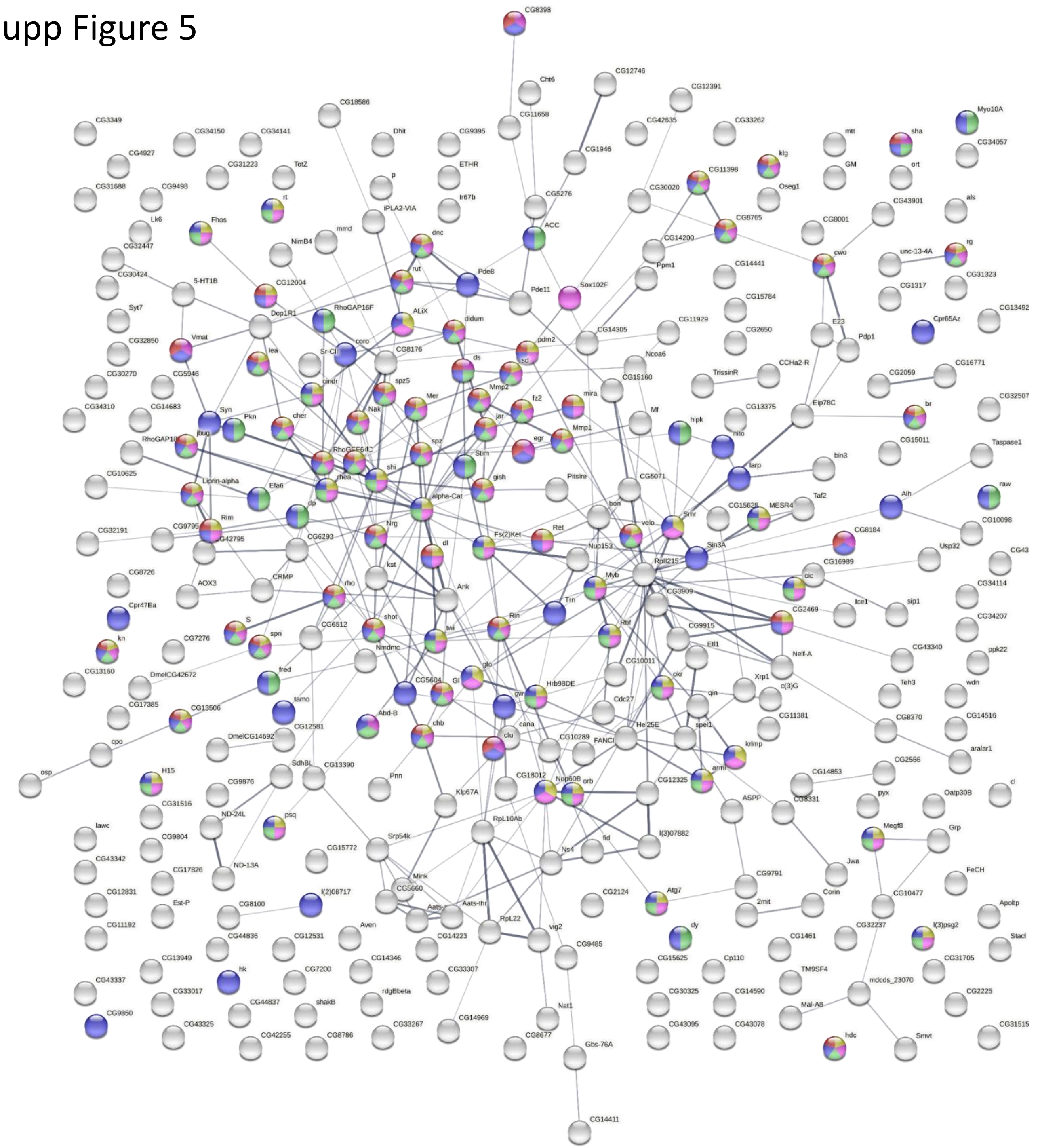

cell development FDR = 2.39e-07

cell differentiation FDR = 9.63e-07

anatomical structure morphogenesis FDR = 1.64e-06

anatomical structure development FDR = 7.46e-06

neurogenesis FDR = 8.61e-06

Supp Figure 6

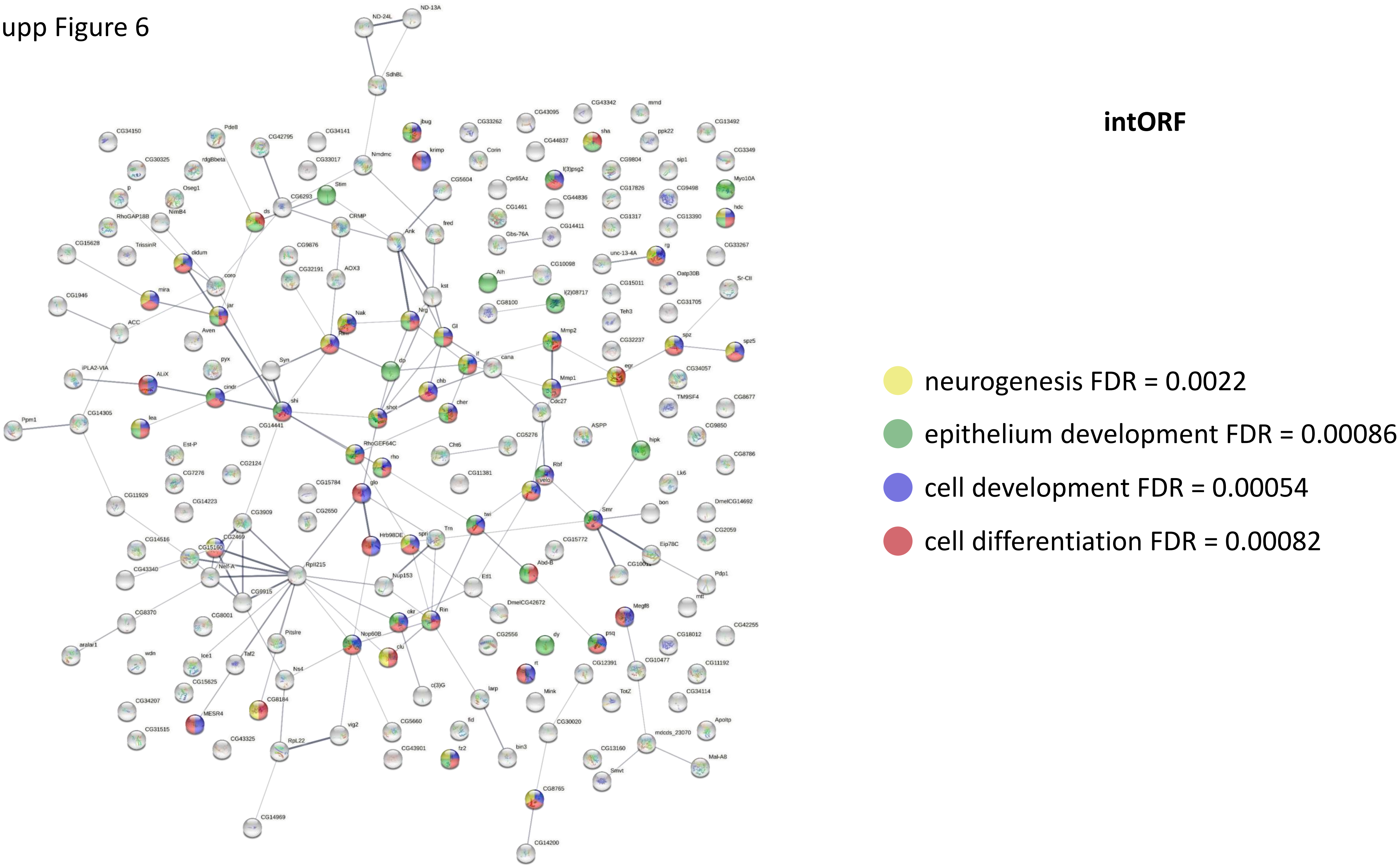

Supp Figure 7

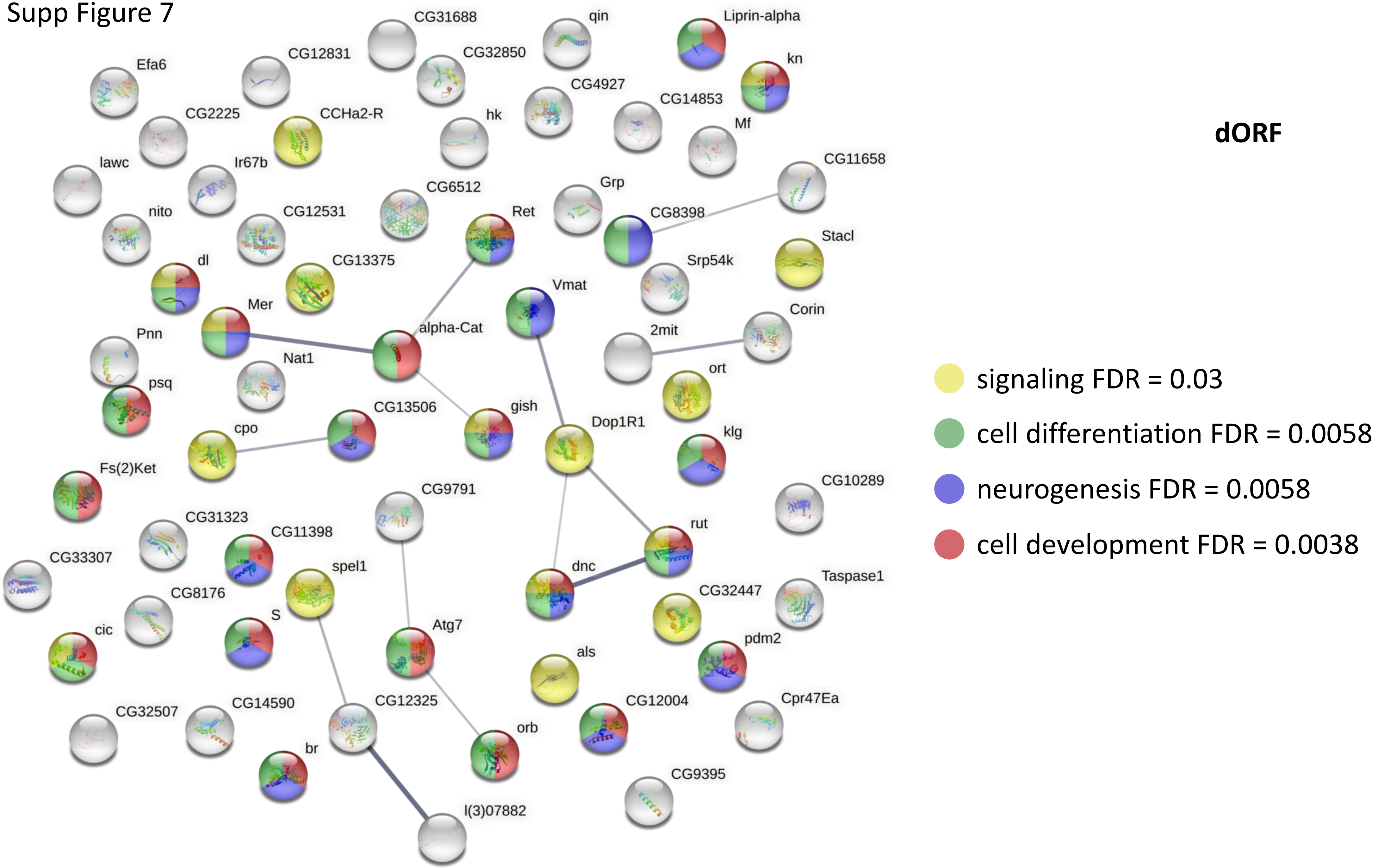

Supp Figure 8

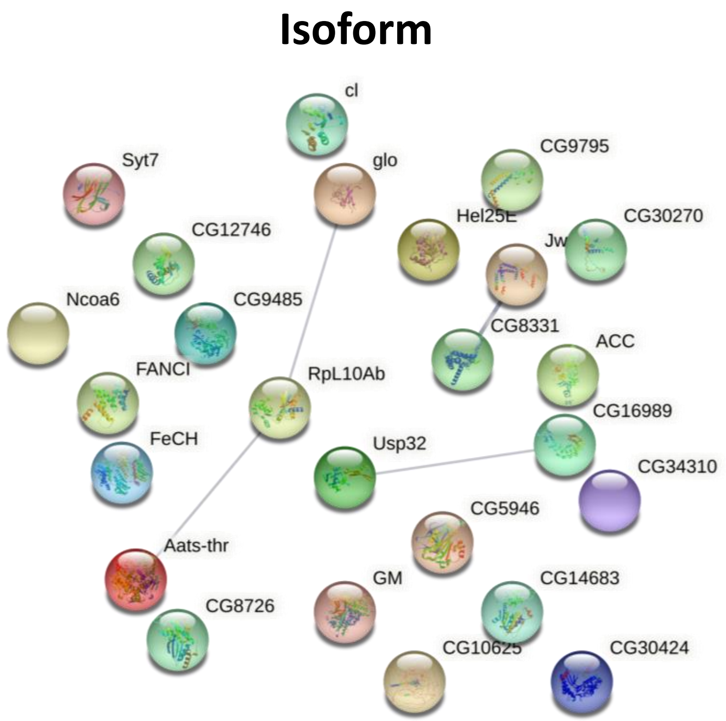


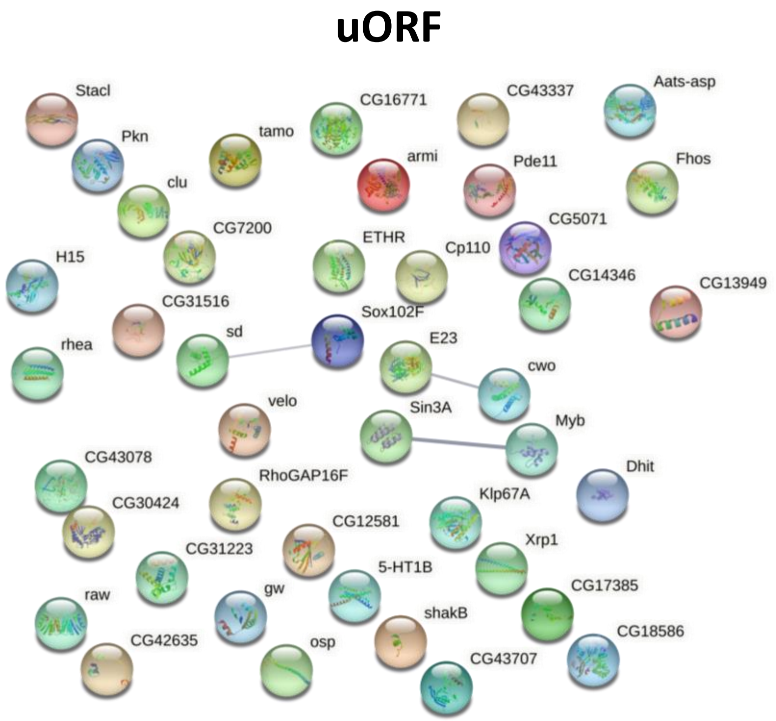

Supp Figure 9

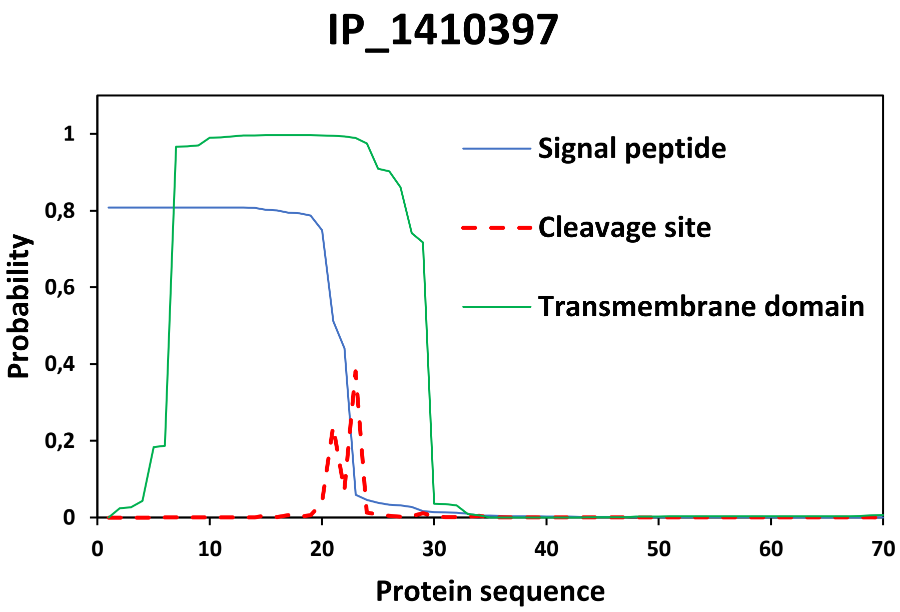

Supp Figure 10

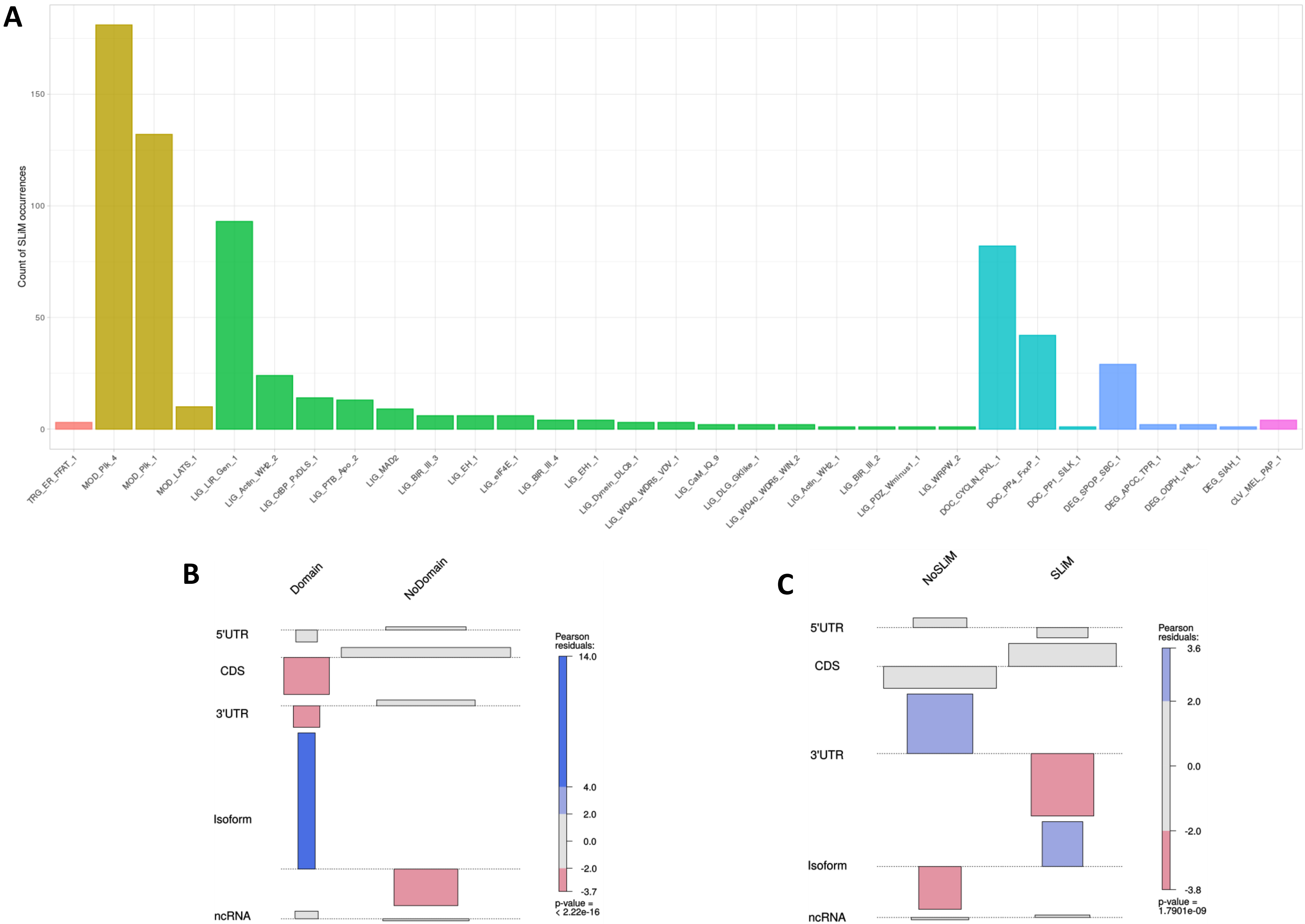

Supp Figure 11

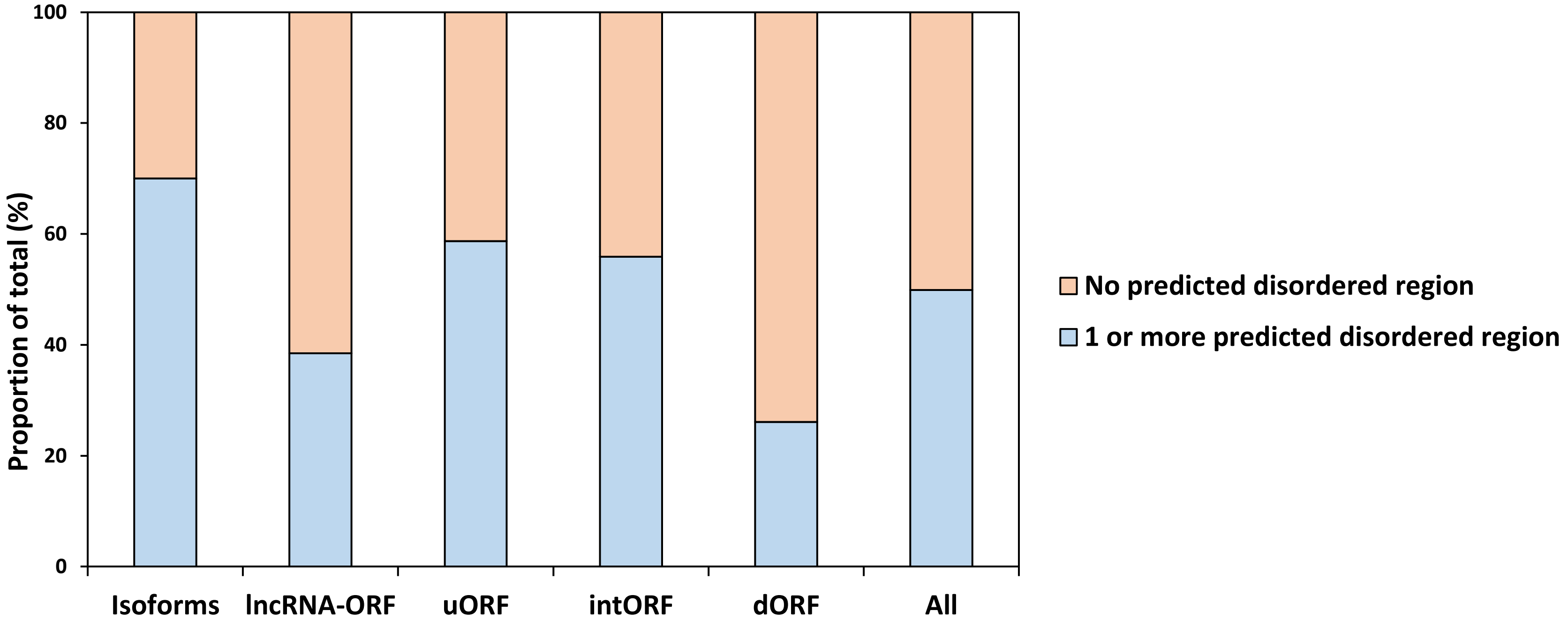

Supp Figure 12

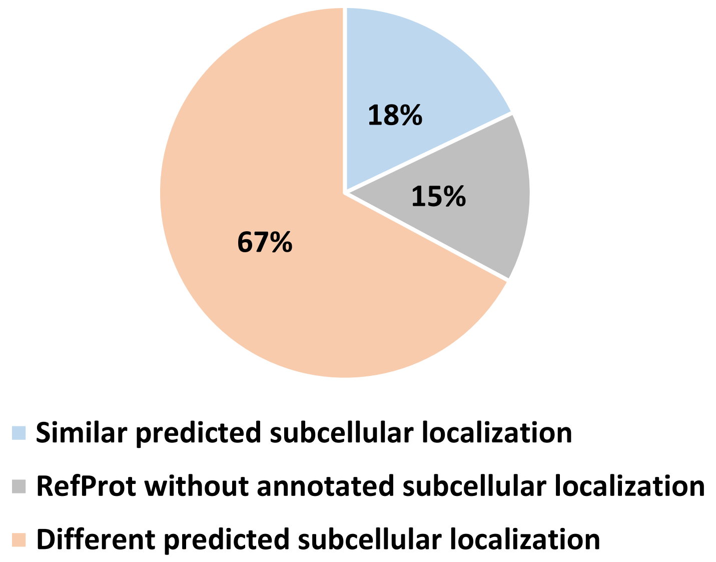

Supp Figure 13

A

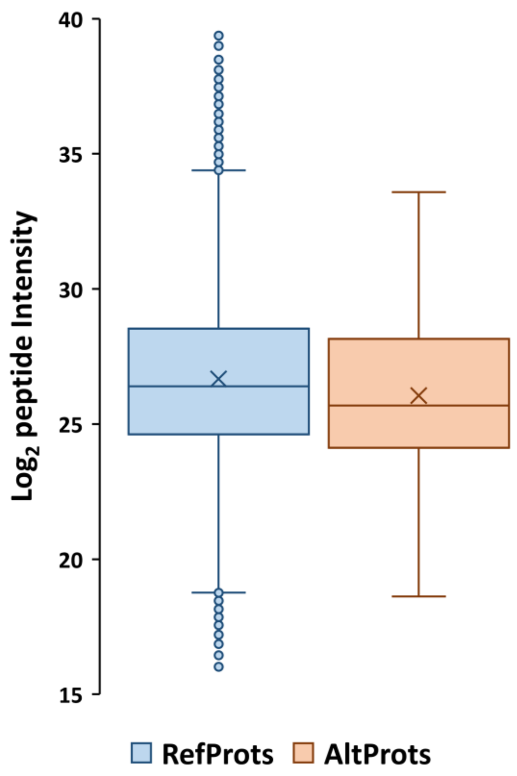


B

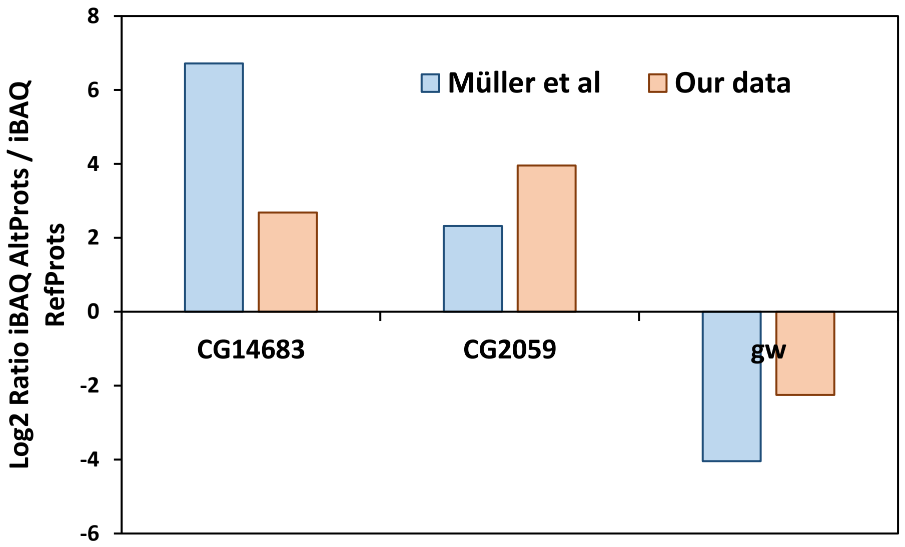

Supp Figure 14

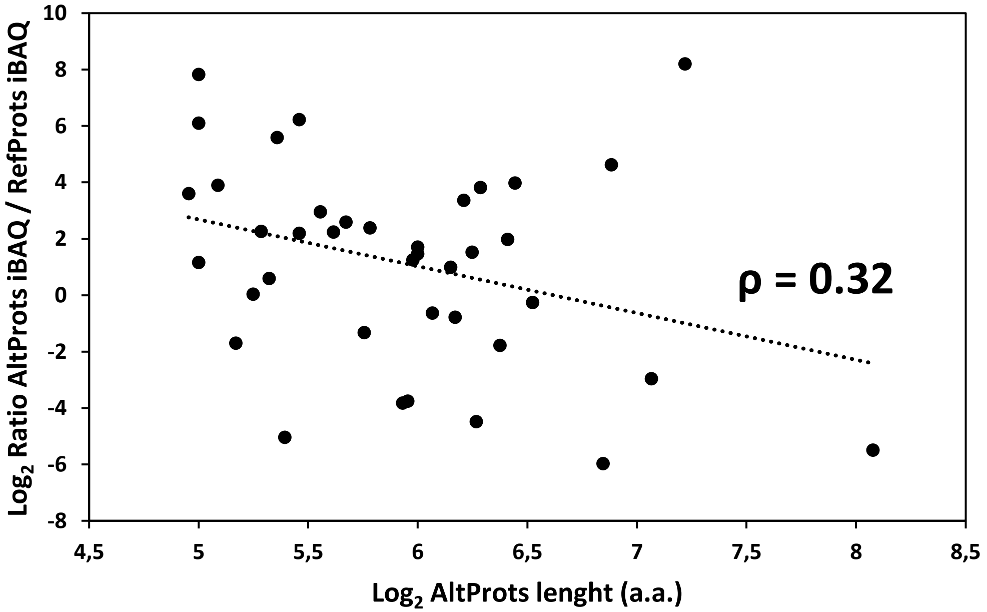

*Supplementary Figure legends:*

**Supplementary Figure 1: Metrics of the different alternative proteins identified in *Drosophila melanogaster*.** A. Andromeda score distribution for peptide spectrum matches of RefProts identified under default MaxQuant settings and AltProts using optimized parameters. B. Distribution of the amino acid length of the AltProts/Isoforms identified in this study. C. Venn diagram representing the overlap of AltProts/Isoforms identified in this study compared to AltProts/Isoforms with MS evidence in OpenProt and the 129 AltProts (out of the 410 small proteins they identified, 281 being already annotated in UniprotKB) identified by Wang *et al.* 2022.

**Supplementary Figure 2: Proportions of the different types of AltProts identified and predicted in *Drosophila melanogaster*.** Distribution of the newly identified proteins depending on their chromosomal location (A), if they are AltPorts or new Isoforms (C), encoded by mRNA or lncRNA (E) and the location of their corresponding ORFs on mRNA (G). The similar distributions were obtained for predicted AltProts/Isoforms from OpenProt (respectively, B, D, F and H).

**Supplementary Figure 3: Distribution of the alternative proteins and isoforms start codon positions ($Log_2$ transformed) depending on the types of ORFs they are produced from.**

**Supplementary Figure 4: Distribution of the amino acid length of the AltProts/Isoforms identified for each type of ORF (A) and normalized by the total protein counts within each group (B).**

**Supplementary Figure 5: Gene Ontology term analysis of the host genes of the alternative proteins and isoforms identified in this study using STRING v11.5.**

**Supplementary Figure 6: Gene Ontology term analysis of the host genes of the alternative proteins identified from intORFs using STRING v11.5.**

**Supplementary Figure 7: Gene Ontology term analysis of the host genes of the alternative proteins identified from dORFs using STRING v11.5.**

**Supplementary Figure 8: Gene Ontology term analysis of the host genes of new proteins isoforms and alternative proteins identified from uORFs using STRING v11.5.**

**Supplementary Figure 9: Probability of the presence of signal peptide and transmembrane domain within the first 70 amino acids of the AltProt IP_1410397 as predicted by SignalP - 5.0 and TMHMM - 2.0, respectively.**

**Supplementary Figure 10: Predicted domains and SLiMs on AltProts.** A. Counts of the different SLiMs motifs in the AltProts identified in this study. B. Association plots showing the dependency between the presence or lack of protein domains for the different types of ORFs the AltProts are produced from. Blue color represents positive association and pink color represents negative association between the presence or absence of protein domains and the type of AltProt. C. Association plots showing the dependency between the presence or lack of SLiMs for the different types of ORFs the AltProts are produced from. Blue color represents positive association and pink color represents negative association between the presence or absence of SLiMs and the type of AltProt.

**Supplementary Figure 11: Distribution of the AltProts with at least one predicted disordered region predicted by IUPred2A and depending on the types of ORFs.**

**Supplementary Figure 12: Proportion of AltProts with predicted subcellular localization similar or different compared to known subcellular localization of their corresponding RefProts.**

**Supplementary Figure 13: A. Comparison of the distribution of peptide intensities ($Log_2$ transformed) measured for RefProts and AltProts in protocol 2 from the material and methods section as well as data from Müller *et al*. B. Graph representing the $Log_2$ ratio between the iBAQ value of AltProt and corresponding RefProts measured in our study and Müller *et al* for the *CG14683*, *CG2059* and *gw* genes.**

**Supplementary Figure 14: Graph representing the correlation (measured using the Pearson correlation coefficient) between the $Log_2$ ratio of the iBAQ value of AltProt and corresponding RefProts and the AltProts amino acid length ($Log_2$ transformed).**
